# Diversity and Genomic Organization of Non-B DNA Motifs in Haplotype-Resolved Human Genome Assemblies

**DOI:** 10.64898/2026.03.05.709836

**Authors:** Alexander Turco, Nadejda B. Boev, Sushant Kumar

## Abstract

Long-read sequencing and telomere-to-telomere genome assemblies now enable the exploration of previously inaccessible repetitive and structurally complex regions of the human genome. Using 130 haplotype-resolved genome assemblies from 65 individuals across diverse populations, we systematically analyzed six major classes of non-B DNA motifs. By evaluating their biophysical stability, we distinguished structurally stable motifs from those forming unstable secondary structures and mapped their distribution across individuals and genomic contexts. Our work revealed significant variation at the population level in both motif abundance and predicted structural stability, uncovering previously unrecognized diversity in non-B DNA landscapes. Non-B DNA motifs exhibit notable, structure-specific enrichment in highly repetitive and evolutionarily dynamic regions that remain largely unresolved in short-read-based genomes, including centromeres, segmental duplications, structural variant breakpoints, and mobile element insertions. Our findings provide a refined view of the potential secondary-structure organization within repetitive regions of the human genome and highlight structural stability as a key factor shaping the distribution of non-B DNA motifs in regions linked to genome instability, evolution, and human variation.

## Introduction

The advent of long-read whole-genome sequencing (lrWGS) techniques such as PacBio HiFi and Oxford Nanopore has enabled the creation of gapless, telomere-to-telomere assemblies. These technological advances have overcome the limitations of bacterial artificial chromosome-based assembly and short-read whole-genome sequencing (srWGS), which struggle to resolve structurally complex regions of the genome (1). The nearly complete and highly accurate assemblies include highly repetitive regions, previously considered “junk” DNA or viewed as by-products of evolutionary processes (2). In particular, the Telomere-to-Telomere (T2T) CHM13 human reference genome represents the first truly complete human genome sequence and adds nearly 200 million base pairs that were unresolved in the previous human reference genome, GRCh38 (1). Building on these technologies, collaborative initiatives such as the Human Genomic Structural Variation Consortium (HGSVC) and the Human Pangenome Reference Consortium (HPRC) have further advanced these efforts by generating high-quality, haplotype-resolved assemblies from individuals representing diverse human ancestries (3, 4, 5, 6, 7).

The availability of these T2T assemblies now enables detailed exploration of repetitive DNA, which comprises roughly 50% of the human genome and is known to drive evolution, induce variation, and regulate critical biological processes such as gene expression (8). These repetitive sequences include non-B DNA (nBDNA) motifs, which have the unique ability to adopt DNA conformations distinct from the canonical right-handed B-form double helix (2, 9). Among the various nBDNA conformations, several well-characterized secondary structures include cruciform/hairpin, H-DNA (triplex), slipped-strand, G4s (tetraplexes), Z-DNA, and bent DNA. These non-B DNA conformations are known to influence various biological processes, including DNA replication, gene regulation, recombination, epigenetic modification, and overall genomic stability (10,11,12,13,14). The nBDNA sequence motifs are not randomly distributed but are preferentially located within regulatory and structurally dynamic regions of the genome. In particular, nBDNA motifs are enriched in highly conserved regions and in genetic instability hotspots, including promoters, replication origins, and telomeres (2). For instance, G-quadruplex (G4) structures are commonly found in promoter regions and telomeric repeats, with nearly 40% of annotated human genes containing one or more promoter G4s (15). Similarly, Z-DNA motifs are enriched within core promoter regions across multiple species (16), while inverted repeats (IRs) capable of forming cruciform structures are concentrated near gene termini, start codons, 5’ untranslated regions, and other transcriptionally active loci (17).

nBDNA motifs are increasingly recognized as contributors to large-scale genomic variations. For instance, prior genomic analyses have shown that potential nBDNA-forming regions of the genome exhibit elevated rates of single-nucleotide polymorphisms (SNPs) and small insertion & deletions (INDELs), suggesting that structures induced by these motifs contribute to increased mutability (18). Additionally, direct repeats (DRs) and mirror repeats (MRs) can generate slipped-strand and H-DNA structures that interfere with DNA polymerases and stall replication forks, thereby promoting fork collapse, double-strand breaks, and larger genomic rearrangements, such as duplications, deletions, and insertions (19, 20). Due to their diverse molecular effects, nBDNA motifs are essential for key cellular processes, including transcriptional regulation and telomere maintenance, yet they can also induce genomic instability (21). For example, putative nBDNA motifs have been found to be enriched at translocation and deletion breakpoints in cancer genomes (22, 23, 24), as well as at copy number variant breakpoints (25). Cruciform-forming IRs have also been shown to increase chromosomal instability in budding yeast (26). Another major source of genomic variation stems from Mobile Element insertions (MEIs), which are closely linked to nBDNA structures (21). nBDNA motifs are highly abundant in transposable elements and may influence the lifecycle of these MEIs, including their integration into the genome (27, 28). G4 motifs are particularly enriched and more stable in evolutionarily younger SVAs (29, 30). Collectively, these findings suggest that nBDNA motifs can contribute to both localized and large-scale genomic instability.

Although several studies have characterized nBDNA motifs in the human genome using computational methods, most have relied on analyses of a single reference assembly or short-read data (22, 31, 32, 33). For example, the Non-B DNA Motif Database provides genome-wide annotations of nBDNA motifs on the GRCh38 reference genome, identified through computational search criteria for each motif type (31). With long-read sequencing, the resolution of repetitive and structurally complex regions has improved the identification of nBDNA motifs among humans, apes, and other T2T assemblies (34, 35, 36). For example, the completion of the T2T-Y chromosome led to the discovery of over 825,000 repetitive sequence motifs predicted to form nBDNA structures (37). Despite these advancements, we still lack a comprehensive understanding of the landscape of nBDNA motifs across diverse human populations and their role in genomic variations. Moreover, although previous reports suggest that approximately 13% of the human genome consists of repetitive motifs capable of forming nBDNA structures based on sequence composition, their stability and functional significance have yet to be fully investigated (10, 34). Many factors, including negative supercoiling of DNA, chromatin structure, and the presence of various binding proteins, also influence the formation and stability of these non-canonical DNA structures (2).

With the continued rise of lrWGS-based accurate T2T genomic assemblies, it is now possible to explore the nBDNA landscape in highly repetitive regions of the human genome and to understand their role in driving genomic variations. Therefore, we employed 130 near-complete, haplotype-specific genome assemblies from 65 diverse individuals in the HGSVC3 study (7) to address this question. These assemblies reveal highly complex genomic regions, allowing for precise annotation of nBDNA motifs within centromeres, segmental duplications (SDs), structural variants (SVs), and mobile element insertions (MEIs). In this study, we primarily focused on six major motif classes, including MRs, IRs, DRs, A-Phased repeats (APRs), G4s, and Z-DNA, which can form into distinct nBDNA secondary structures with unique topological and biological properties. In particular, we systematically examined their genome-wide distribution, population-specific variation, and enrichment in regions linked to structural complexity and genome instability. While previous studies have primarily explored the relationship between nBDNA motifs and shorter genomic variants, such as SNPs and INDELs, our haplotype-resolved framework enables investigation of these motifs in the context of longer SVs, SDs, and MEIs. This large-scale, haplotype-resolved study provides a comprehensive view of the nBDNA motif landscape in complete human genomes and improves our understanding of how non-canonical DNA secondary structures affect genetic diversity and genome stability.

## Results

### Genome-wide Profiling of Non-B DNA Motifs Across 65 Telomere-to-Telomere Human Assemblies

We characterized the genome-wide nBDNA landscape of haplotype-specific assemblies for 65 diverse human samples (130 haplotypes) from the HGSVC3 study. Briefly, these samples were sequenced using Hi-Fi- and ONT-based long-read technologies, assembled with the Verkko algorithm, and aligned to both CHM13v2.0 and GRCh38.p14 reference genomes (7). To identify nBDNA structures, we split haplotype-aligned BAM files by chromosome. We annotated G4 motifs using the Quadron tool (38) and retained those with a stability score greater than 19. Concurrently, we annotated MRs, DRs, IRs, APRs, and Z-DNA motifs using the non-B gfa tool (31) (**Figure 1A**). Overall, we observed that cumulative nBDNA motif coverage was slightly higher in CHM13-aligned assemblies than in GRCh38, with this difference consistently observed across both haplotypes and autosomes and sex chromosomes (**Figure 1B)**. The variation in nBDNA coverage among haplotypes aligned to different human genome reference builds likely reflects the greater completeness of CHM13 compared to GRCh38. However, the genome-wide coverage for various motifs was consistent between the references (**Figure 1C**). For example, IRs (4.5%), MRs (2.7%), and DRs (2.3%) were most frequent across all samples, while APR (0.4%), G4+ (0.23%), G4- (0.23%), and Z-DNA (0.21%) motifs each made up less than 1% of the human genome.

**Figure 1.**
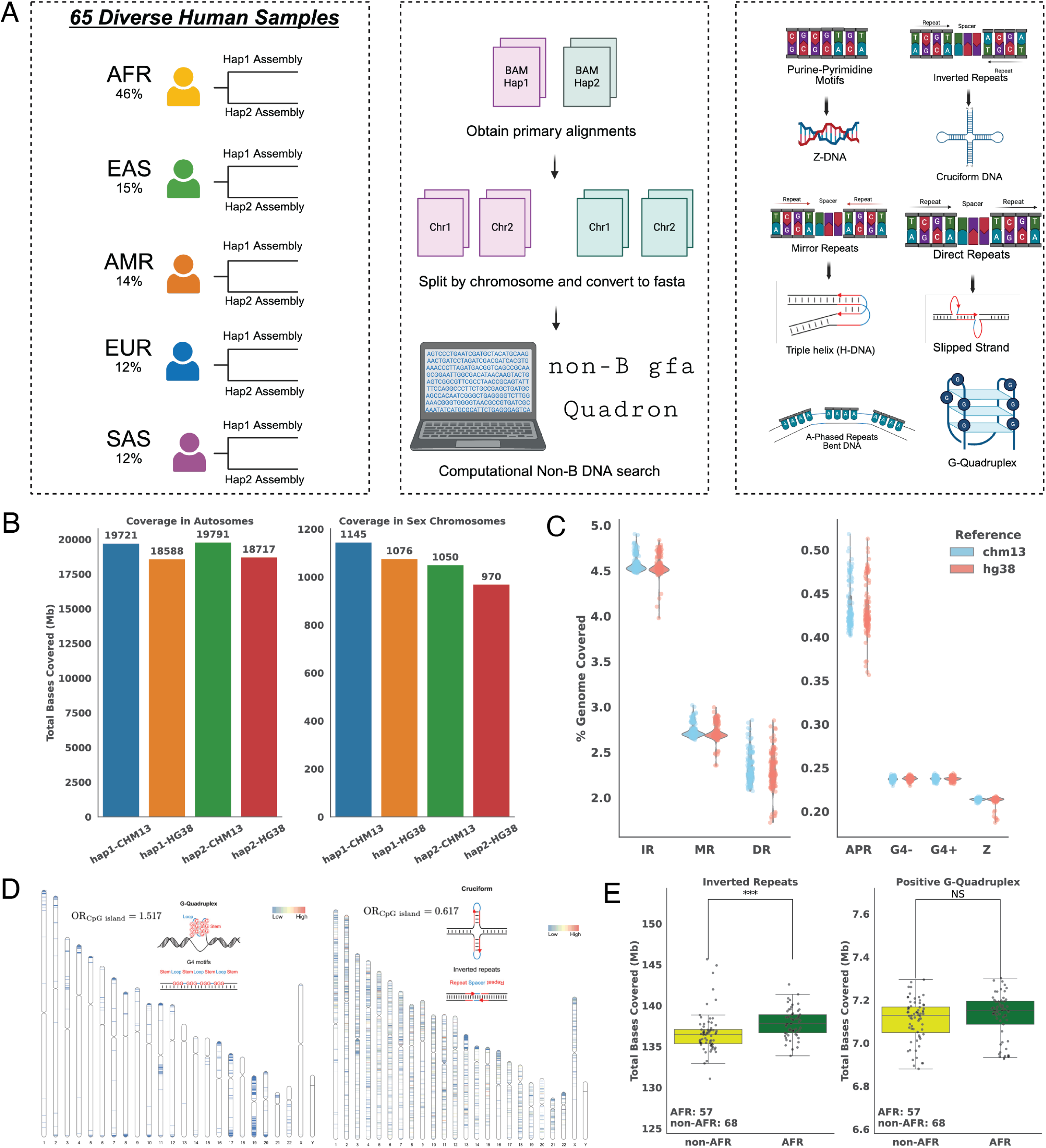
Overview of non-B DNA annotation workflow across 130 diverse human haplotypes. **(A)** Cohort composition of 65 diploid individuals spanning five continental superpopulations (African, East Asian, American, European, South Asian), with haplotype-resolved assemblies generated for each individual by the HGSVC. Haplotype-specific BAMs aligned to GRCh38 and T2T-CHM13v2.0 were split by chromosome and converted to FASTA for computational non-B DNA motif annotation. Two tools were applied: non-B gfa for broad motif classes and Quadron for G-Quadruplexes. Non-B DNA motif classes include Z-DNA, inverted repeats (cruciform DNA), mirror repeats (H-DNA), direct repeats, A-phased repeats, and G-Quadruplexes. **(B)** Cumulative non-B DNA coverage across haplotypes in autosomes and sex chromosomes, stratified by reference genome (GRCh38 vs T2T-CHM13v2.0) and haplotype (1 vs 2). Coverage is reported as the summed base-pair span of annotated motifs across all haplotypes. **(C)** Percentage of genome covered by distinct non-B DNA classes in haplotype-resolved assemblies aligned to T2T-CHM13v2.0 (blue) and GRCh38 (red). Motif types include inverted repeats (IR), mirror repeats (MR), direct repeats (DR), A-phased repeats (APR), Z-DNA (Z), and G-quadruplexes (G4). **(D)** Chromosomal ideograms of motifs recurrently annotated in five or more haplotypes aligned to CHM13. Odds ratios for G4s and IRs in CpG islands highlight the enrichment of G4s in these regions (1.517) and depletion of IRs in these regions (0.617). **(E)** Population-level comparisons of non-B DNA coverage. Per-haplotype base-pair coverage of inverted repeats (IRs) is significantly higher in African (AFR) haplotypes compared to non-African Haplotypes (Mann-Whitney U test, p < 0.001), while no significant difference is observed for positive-strand G-quadruplexes (NS).

Upon analyzing the chromosomal distribution of shared G4 and IR motifs across haplotypes (CHM13-aligned), we observed distinct localization patterns for the two motif types **(Figure 1D)**. For example, G4s present in five or more haplotype assemblies tend to occur preferentially in telomeres, gene-rich regions, and are highly enriched in CpG islands (Odds Ratio (OR) = 1.517, p < 0.05, Fisher’s exact test), aligning with their role in transcriptional regulation. In contrast, IRs shared among five or more haplotype assemblies were distributed more uniformly across the genome and were depleted from CpG islands (OR = 0.617, p < 0.05). Next, we examined associations between the distribution of nBDNA motifs and population groups. Overall, haplotypes from the African (AFR) population group exhibited significantly higher IR base coverage than those from non-African (non-AFR) individuals (p < 0.001, Mann-Whitney U test; **Figure 1E**). However, G4 motif coverage was similar between AFR and non-AFR haplotypes (p = NS). These observations might suggest that IRs potentially contribute to the elevated levels of genomic variation observed in the AFR population. In contrast, G4s maintains similar distributions across populations because of its presence in critical genomic functional regions.

### Differential Stability and Functional Enrichment of Inverted Repeats and G-Quadruplex Motifs Across the Human Genome

nBDNA motifs vary significantly in their likelihood to form secondary structures in vivo. While many sequences display nBDNA motif patterns, not all will form stable or biologically active secondary structures due to differences in the cellular environment, chromatin state, and the stability or folding energy of the underlying sequence (2). Therefore, besides mapping the genome-wide distribution of nBDNA motifs, we analyzed the sequence features that influence their stability and whether this stability affects their localization within specific genomic elements. We used Quadron-predicted stability scores (Q) for G4 motifs and seqfold-based free-energy predictions for IR sequences (see Methods for more details) to assess their structural stability. The sequence architecture of IR motifs comprises two complementary repeat arms that form double-stranded stems, separated by a spacer sequence that forms a loop (**Figure 2A**).

**Figure 2.**
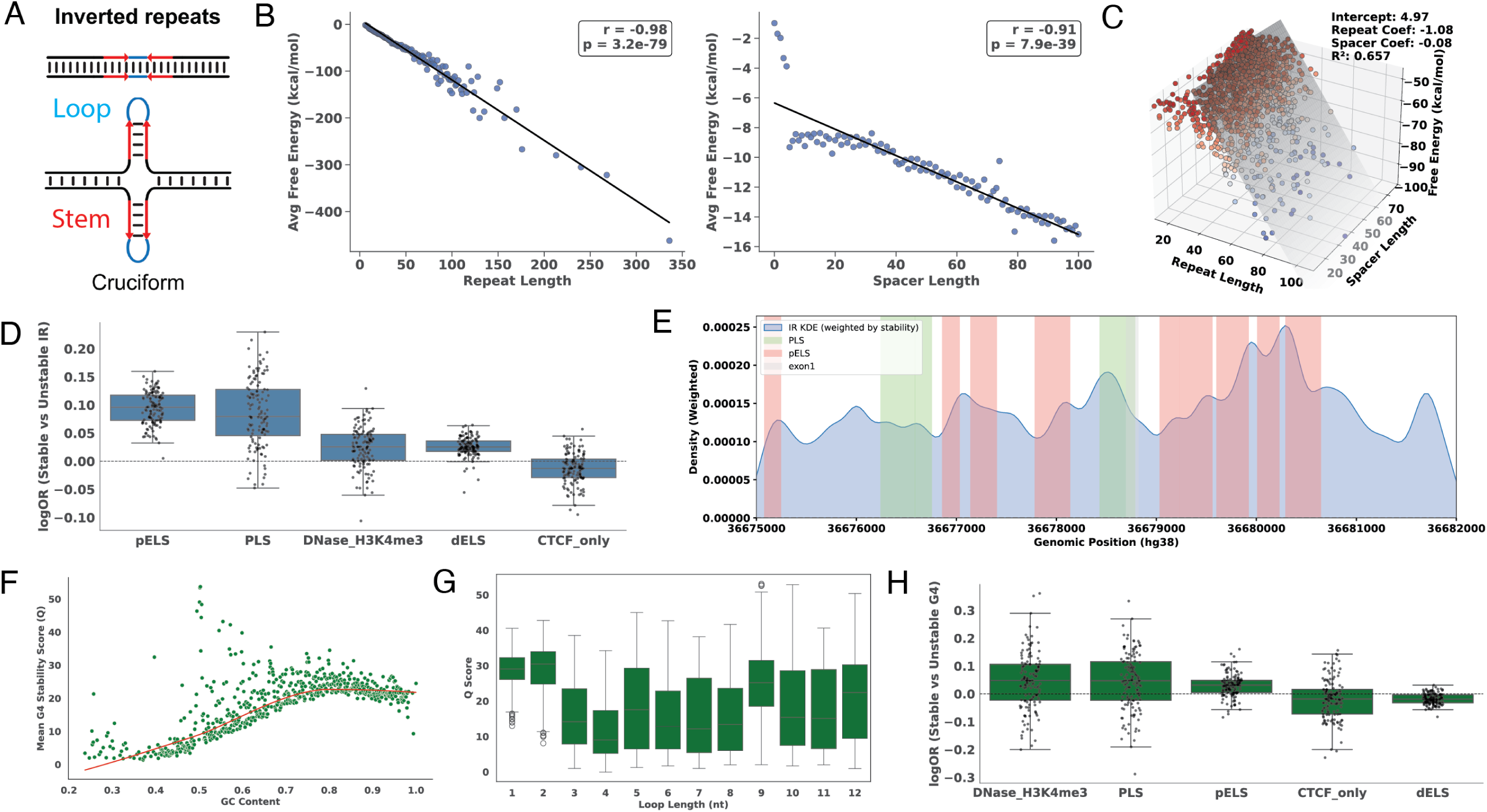
Stability determinants and regulatory enrichment of inverted repeats (IR) and G-quadruplexes (G4). **(A)** Inverted repeat motifs consist of repetitive complementary sequences (stem) separated by a nonrepetitive spacer (loop) and can promote the formation of cruciform or hairpin structures. **(B)** Relationship between inverted repeat (IR) free energy and sequence architecture in a single haplotype assembly. Average free energy decreases with increasing repeat length (left) and spacer length (right). **(C)** Multivariable ordinary least squares (OLS) regression of free energy across all haplotypes, incorporating both repeat and spacer length, demonstrates that each contributes significantly to IR stability (R2=0.657). **(D)** Log odds ratios (Fisher’s exact test) of stable versus unstable IRs intersecting with candidate cis-regulatory elements (cCREs). **(E)** Kernel density estimate (KDE) of IR positions across haplotypes within the CDKN1A locus, weighted by stability, reveals enrichment of stable cruciform in specific cCREs. **(F)** G-quadruplex (G4) stability scores as a function of GC content, showing a positive association that plateaus beyond ∼0.80 GC fraction. **(G)** Relationship between G4 loop length and stability, with shorter loops associated with higher Q-scores and longer loops with reduced stability. **(H)** Log odds ratios of stable versus unstable G4s intersecting with cCREs, analogous to the IR analysis, highlighting regulatory enrichment patterns.

The potential of an IR to form a cruciform structure mainly depends on the lengths of these repeats and spacers. Therefore, we included all predicted IRs with a minimum repeat length of six nucleotides and a maximum spacer length of one hundred nucleotides. We observed a strong and significant negative correlation between repeat arm length and average free energy (r = −0.98, p-value = 3.2E-79), indicating IRs with longer repeat arms tend to be more stable (**Figure 2B**). Surprisingly, a similarly strong negative correlation was found between spacer length and average free energy (r = −0.91, p=7.9E-39), suggesting that even longer spacers can contribute to increased sequence stability (**Figure 2B**). This contrasts existing experimental models of cruciform formation, which predict reduced stability for extended loop regions (39, 40, 41); however, folding free energies inherently become more negative with increasing sequence length. Moreover, long IRs remain relatively understudied experimentally, and recent work has begun to examine how longer IRs might still attain stable cruciform structures (42). To assess the relative contributions of these sequence features, we used a multiple linear regression model including repeat and spacer length, which explained 65.7% of the variance in folding free energy (R² = 0.657; Figure 2C). Repeat length showed a stronger independent association with free energy (β = −1.08; standardized β = −2.66) than spacer length (β = −0.077; standardized β = −1.44), with both predictors contributing significantly (p < 0.001).

After examining the sequence determinants of IR stability, we then asked whether structurally stable IRs tend to localize to candidate cis-regulatory regions (cCREs) (43). Overall, without stratifying by stability, IRs did not show strong enrichment across cCRE categories relative to the rest of the genome (median logOR = 0; S1 Fig. **1A**), suggesting that IRs are not generally concentrated in regulatory regions. Next, we evaluated whether structural stability, rather than IR presence alone, affects regulatory localization. Using a stability threshold of 0 kcal/mol, we compared stable and unstable IRs overlapping cCREs within each haplotype. We found that stable IRs were consistently more enriched than unstable IRs across multiple cCRE classes, especially in enhancer (pELS) (median logOR = 0.10) and promoter (PLS) elements (median logOR = 0.08) (**Figure 2D, Supplement Table 10**). To further support the functional significance of IR stability, we examined the CDKN1A (p21) locus, which contains previously characterized cruciform-forming IRs that help facilitate p53 binding (44). Consistent with our results, we observed stable IRs clustered within promoter and enhancer regions at this locus (**Figure 2E).**

To evaluate whether similar sequence-stability relationships apply to other nBDNA motifs, we analyzed G4 stability. We observed that G4 stability increased with GC content but plateaued beyond approximately 75% GC content (**Figure 2F**). Next, we investigated the relationship between the G4 loop length and its stability score. We found that G4 motifs with shorter loop lengths (1-2 nucleotides) had higher stability scores than those with longer loops (**Figure 2G)**. Interestingly, loop lengths of 9 and 12 nucleotides confer greater stability, suggesting that the relationship between loop length and stability may be nonlinear and that specific configurations support stable G4 formation. Finally, to determine whether G4 stability itself is associated with regulatory elements, we compared G4 stability across cCREs. Both stable (Q > 19) and unstable (Q < 19) G4s were globally enriched in promoter-like (PLS) and proximal enhancer-like (pELS) regions. However, when directly comparing stable and unstable G4s within haplotypes, we observed very small differences in relative enrichment across cCRE classes (median logOR ≍ 0; **Figure 2H**). Stable G4s showed slightly higher enrichment in DNAse-H3K4me3 and promoter-associated elements, but minimal separation overall. To evaluate the broader regulatory landscape, we also assessed cCRE enrichment across other nBDNA motif types (**S1 FIG 1A**). Notably, Z-DNA motifs exhibited modest but consistent enrichment across multiple cCRE categories, while DRs and APRs tended to be depleted across cCREs.

### Centromeric Regions Exhibit High Density and Distinct Organization of Non-B DNA Motifs

These long-read-based, haplotype-specific assemblies also enabled us to explore various highly repetitive regions that are functionally critical, such as centromeres. Therefore, we examined the prevalence and organization of nBDNA motifs in completely and accurately assembled centromeric DNA and within the active alpha satellite (ASAT) higher-order repeat (HOR) arrays. Across all haplotypes and chromosomes, IRs accounted for approximately 3-8% of the total centromeric bases, with DRs and MRs also constituting notable proportions (**Figure 3A**). The variation observed across haplotypes likely results from differences in centromere length and HOR array organization, as centromeric domains vary substantially among individuals (7, 45). Within HOR arrays across all chromosomes, IRs remained the most abundant motif class, comprising 2-8% of active array DNA, followed by DRs and MRs (**Figure 3A**). Although we detected stable G4s within centromeric regions, their frequency and coverage were markedly reduced across both the broader centromeric domains and within active ASAT HOR arrays (**S2 FIG A**), a finding consistent with previous work describing the underrepresentation of G4 motifs at centromeres when compared to other nBDNA motif types (36).

**Figure 3.**
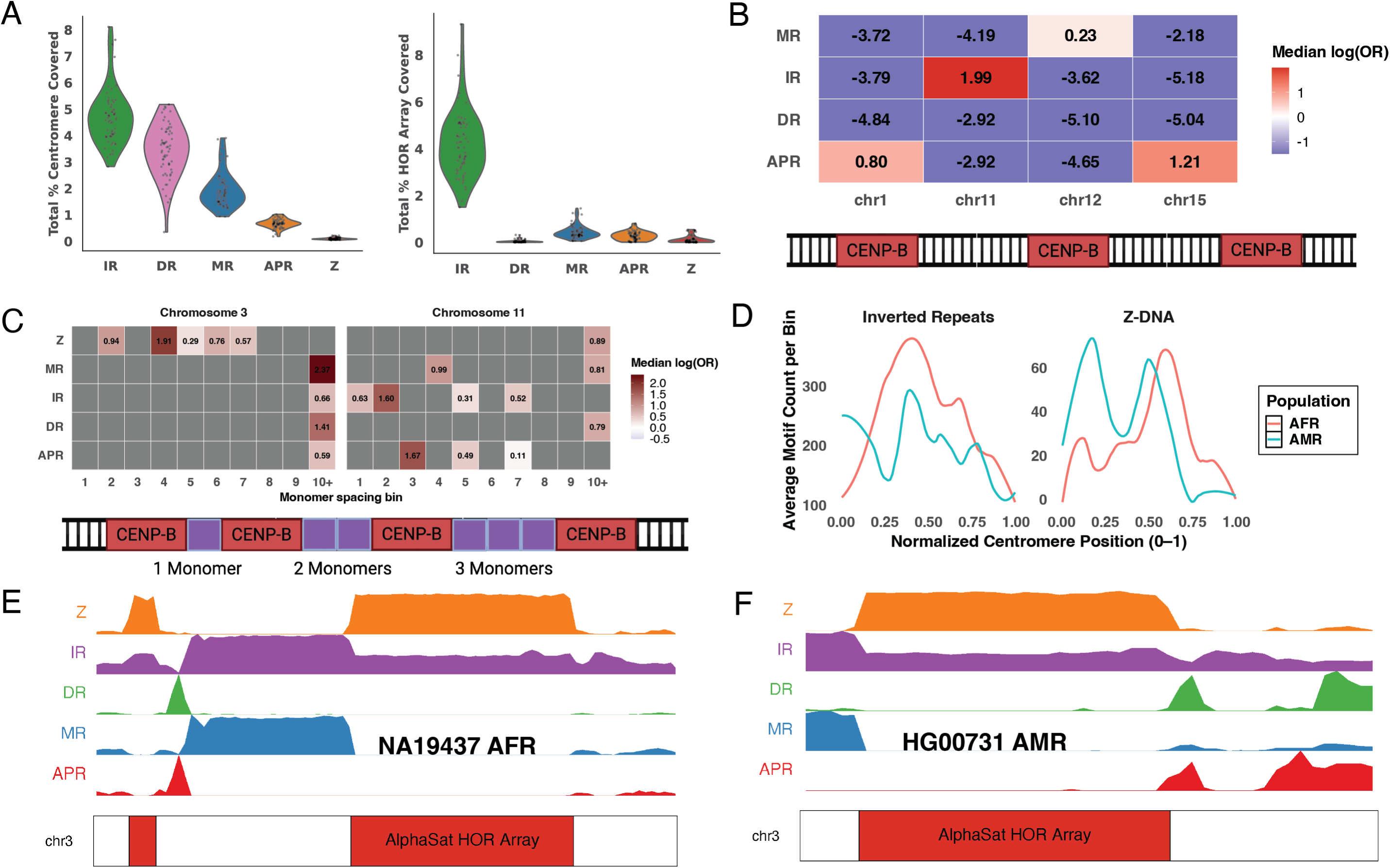
Non-B DNA motif density, enrichment, and organization within fully assembled human centromeres. **(A)** Fraction of total centromeric sequence (left) and active α-satellite higher-order repeat (HOR) arrays (right) covered by major non-B DNA motif classes across haplotypes. Inverted repeats (IRs) represent the largest fraction of centromeric and HOR-array sequence, followed by direct repeats (DRs) and mirror repeats (MRs), whereas G4 and Z-DNA motifs are comparatively rare. **(B)** Median log odds ratios of non-B DNA motif enrichment within CENP-B boxes across haplotypes for chromosomes 1, 11, 12, and 15. Chromosome-specific enrichments include IRs in CENP-B boxes on chromosome 11, and A-phased repeats (APRs) in CENP-B boxes on chromosomes 1 and 15 **(C)** Enrichment of non-B DNA motifs across CENP-B monomer-spacing categories (1-10+ monomers) for representative centromeres on chromosomes 3 and 11. Certain motifs display spacing-dependent enrichments, including Z-DNA on chromosome 3 and APRs on chromosome 11. **(D)** Normalized density profiles of inverted repeats (IRs) and Z-DNA motifs across chromosome 3 centromeres (scaled position 0-1), separated by population group. Both African (AFR) and American (AMR) haplotypes show peaks aligned with α-satellite HOR arrays, though with notable differences in positional organization. **(E-F)** Representative chromosome 3 centromeres from samples NA19437 (AFR) and HG00731 (AMR), showing motif-density tracks for Z-DNA, IRs, DRs, MRs, and APRs across the α-satellite HOR array.

Furthermore, we annotated CENP-B boxes within centromeric DNA and performed motif-enrichment analyses for each haplotype assembly using the Genomic Centromere Profiling workflow (46). Several non-B DNA motifs showed chromosome-specific enrichments, including IRs within CENP-B boxes on chromosome 11 and APR motifs within CENP-B boxes on chromosomes 1 and 15 (median logOR > 0; Figure **3B**). Genome-wide enrichment profiles for CENP-B box-associated nBDNA motifs across all chromosomes showed enrichment was largely absent outside chromosomes 1, 11, 12, and 15 (**S3 FIG A**). Additionally, we examined the organization of ASAT arrays based on the spacing of monomers between consecutive CENP-B boxes, thereby allowing us to assess nBDNA within these intervals. Several motifs exhibited enrichment patterns at specific monomer intervals across chromosomes, including Z-DNA in chromosome 3 and APRs in chromosome 11 (median logOR > 0; **Figure 3C**). The genome-wide CENP-B box monomer spacing enrichment across all chromosomes indicated that while spacing-dependent patterns are detectable on specific chromosomes, they are not universally observed **(Supplementary Figure S3B)**.

Centromeric sequences are highly variable, even among individuals within the same superpopulation, prompting us to examine whether their distribution exhibits population-specific patterns. IR and Z-DNA motifs displayed distinct peaks at specific normalized positions corresponding to the ASAT arrays (**Figure 3D**). Although the underlying centromeric sequences and organization vary significantly in the human population, the nBDNA motifs within the ASAT arrays were consistently present across all haplotypes. To directly visualize these relationships, we generated motif density maps in centromeres on chromosome 3 for representative haplotypes from African (NA19437) and American (HG00731) individuals (**Figures 3E-F**). Within the centromeres of chromosome 3, both haplotypes exhibited a high density of Z-DNA motifs in ASAT HOR arrays; however, the organization and orientation of these arrays differed between individuals, as observed across all haplotypes (**Figure 3D**).

### Non-B DNA Motifs are Enriched in Segmental Duplications

Next, we assessed the distribution of nBDNA motifs within highly repetitive segmental duplications (SDs), which are known hotspots for genomic instability and may play a critical role in the emergence of structural variation (47, 48, 49, 50). We considered SDs that are fixed (shared) across all haplotype-specific genome assemblies in HGSVC3 and the CHM13 reference genome. We categorized these shared SDs as intra- or inter-chromosomal based on whether they occurred within the same or different chromosomes, respectively. Subsequently, we quantified nBDNA motif enrichment separately in intra- and inter-chromosomal SD regions to determine differential nBDNA motif composition. Across the genome, we found that both SD types were enriched for specific nBDNA motifs, though the magnitude of enrichment differed between the two SD classes, with significant median log-odds-ratio differences across haplotypes (**Figure 4A**). The strongest enrichment was observed for DRs, which had the highest odds ratios in both SD types(*x̃_logOR_inter_*= 0.71, *x̃_logOR_intra_*= 0.39). While, G4 motifs (*x̃_logOR_inter_* = 0.16, *x̃_logOR_intra_* = 0.24) and APR motifs (*x̃_logOR_inter_* = 0.22, *x̃_logOR_intra_* = 0.07) also exhibited consistent but lower enrichment than DRs in both SD categories. Interestingly, enrichment patterns differed between intra- and inter-chromosomal SDs, as reflected by shifts in log odds ratios across haplotypes, potentially suggesting that duplication context influences the degree of nBDNA motif enrichment. In contrast, Z-DNA, IRs, and MRs showed little to no enrichment in either SD type, with log odds ratios close to 0. Additionally, we compared enrichment of stable and unstable G4 and IRs to determine whether motif stability is associated with SD localization. Across haplotypes, unstable G4s were slightly more enriched than stable G4s within both intra- and inter-chromosomal SDs (**Figure 4B**). In contrast, IRs were not enriched genome-wide within SDs; however, stable IRs showed higher enrichment than unstable IRs across both SD types, with log odds ratios consistently above zero (**Figure 4C**). The enrichment pattern based on the stability of G4s and IRs also varied widely across haplotypes, with some individuals showing strong enrichment and others showing minimal or no signal. This variability indicates that the relationship between nBDNA motif stability and SD type, and their overlap, differs among haplotypes, potentially reflecting distinct sequence architectures and population-specific differences.

**Figure 4.**
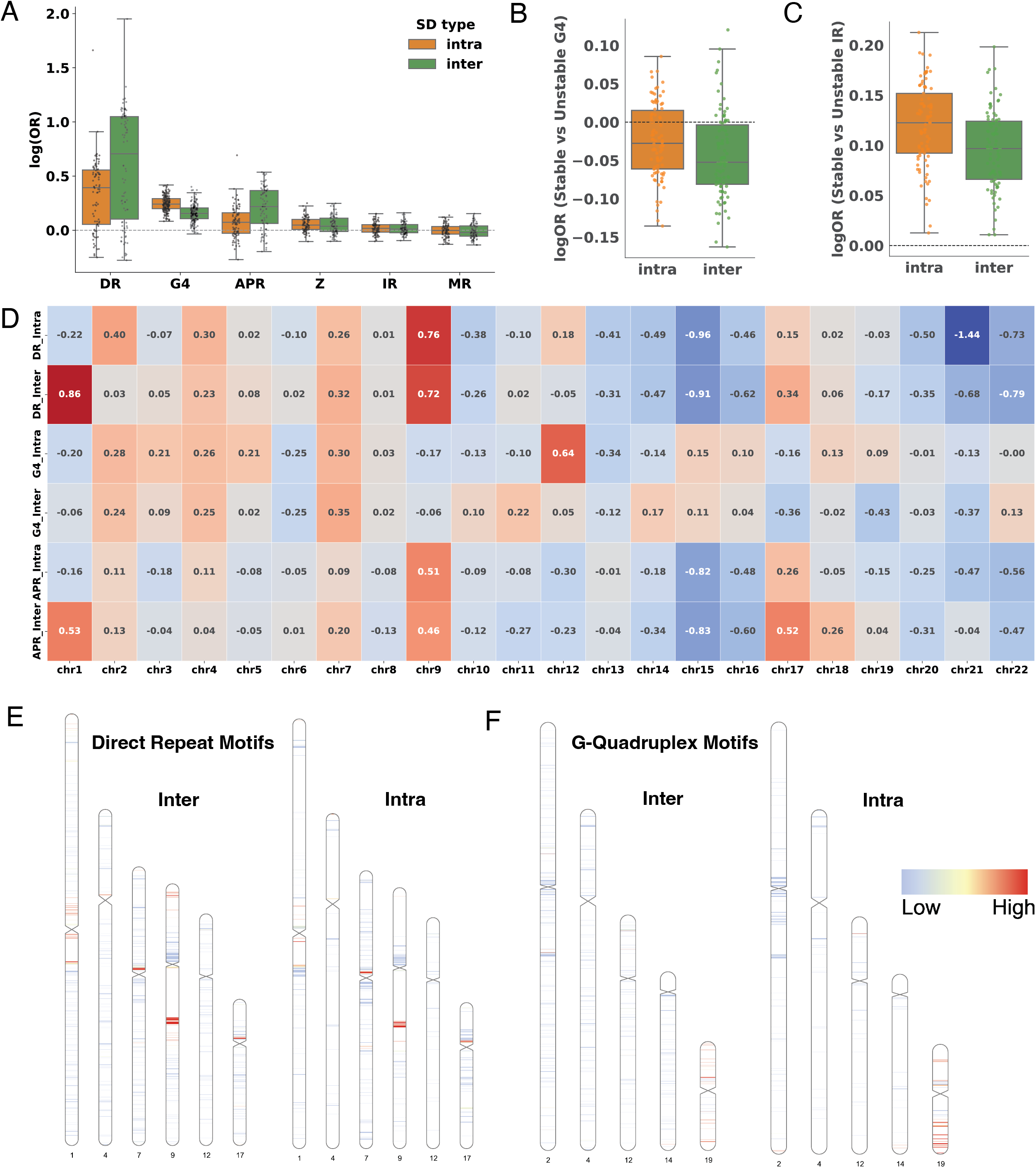
Non-B DNA motifs show distinct enrichment patterns within fixed human segmental duplications (SDs) across 130 phased haplotypes. **(A)** Genome-wide enrichment of non-B DNA motif classes within intrachromosomal and inter-chromosomal SDs relative to the genome background. Direct repeats (DRs) exhibit the strongest enrichment in both SD types, followed by G4 and APR motifs, with different enrichment distributions between intra- and inter-chromosomal duplications. Z-DNA, inverted repeats (IRs), and mirror repeats (MRs) show no enrichment (log odds ratio = 0, Fisher’s exact test). **(B)** Stability-stratified enrichment of G-quadruplex (G4) motifs within SDs, showing that both stable and unstable G4s are enriched relative to genome background, with unstable G4s displaying slightly higher enrichment in both SD types. **(C)** Stability-stratified enrichment of inverted repeat motifs within SDs. Although IRs as a class are not strongly enriched within SDs (log odds ratios close to 0), stable IRs consistently show higher odds ratios than unstable IRs in both intra- and inter-chromosomal SDs, indicating a relative preference for more stable IR configurations within duplicated regions. **(D)** Chromosome-resolved enrichment of DR, G4, and APR motifs within fixed SDs. Inter-chromosomal SDs show strong DR and APR enrichment on chromosomes 1, 9, and 17, whereas intrachromosomal SDs show elevated G4 enrichment on chromosome 12 and along chromosome 19. **(E-F)** Representative density ideograms illustrating spatial distributions of enriched motifs within SDs. DRs are prominent in pericentromeric inter-chromosomal SDs on chromosomes 1 and 9, while G4s show higher density within intrachromosomal SDs on the short arm of chromosome 12 and across gene-rich regions of chromosome 19.

To further characterize this heterogeneity, we stratified stable versus unstable motif enrichment by population groups. Across both intra- and inter-chromosomal SDs, G4 and IR enrichment exhibited marked differences among superpopulations (**S5 Fig A-F**). While overall enrichment patterns were generally consistent across populations, greater variability was observed among haplotypes within African, European, and South Asian groups. We also observed distinct motif-specific patterns of nBDNA enrichment within fixed SDs after chromosome-based stratification (**Figure 4D**). Focusing on the three motif classes most enriched within SDs (DRs, G4s, APRs), we observed apparent chromosome-specific differences between intra- and inter-chromosomal SDs. For example, DR motifs showed strong enrichment, particularly in inter-chromosomal SDs, with prominent signals on chromosomes 1 (*x̃_logOR_inter_* = 0.86), 9 (*x̃_logOR_inter_* = 0.72), and 17 (*x̃_logOR_inter_* = 0.34). G4 motifs displayed high intra-chromosomal SD enrichment on chromosome 12 (*x̃_logOR_intra_* = 0.64), and inter-chromosomal SD enrichment on chromosome 14 (*x̃_logOR_inter_* = 0.17), whereas APRs were enriched within inter-chromosomal SDs on chromosomes 1 (*x̃_logOR_inter_* = 0.53) and 17 (*x̃_logOR_inter_* =0.52). Overall, DR and APR enrichment was stronger within inter-chromosomal SDs, consistent with the observation that these SDs are more frequently found in repeat-dense sub-telomeric and pericentromeric regions of the human genome (51). For DRs, the strongest enrichment was observed in the pericentromeric regions of chromosomes 1 and 9, particularly within inter-chromosomal SDs (**Figure 4E)**. While G4s were highly enriched among intra-chromosomal SDs within the short arm of chromosome 12 and along chromosome 19, which is the most gene-rich chromosome in the human genome (**Figure 4F)**. Together, these trends reinforce that nBDNA motif enrichment within fixed human SDs is not uniformly distributed but rather reflects the local structure and compositional context of the underlying duplicated regions.

### Non-B DNA Motifs are Preferentially Localized near Structural Variant Breakpoints

nBDNA motifs, including G4s and IR, are often associated with the introduction of genomic variations, such as SVs in the human genome (52). Therefore, we analyzed the distribution and stability of all nBDNA motifs in the 2kb flanking regions (up- and downstream) of SVs. Across all haplotypes, we observed a high density of nBDNA motifs proximal to SV breakpoints (**Figure 5A**). MRs and DRs showed narrow density peaks at the breakpoints, while IRs showed a depleted peak, despite covering a significantly larger part of the genome. Interestingly, APRs did not show a similar density pattern around either large insertions or deletions. Next, we examined the stability profiles of IRs and G4s around SV flanking regions to determine whether proximity to SV breakpoints correlates with changes in structural stability. For insertions, IRs displayed increasingly negative free energies (−4 to −12 kcal/mol), indicating greater stability near the breakpoint, followed by a sharp local destabilization (**Figure 5B**). In contrast, G4 stability steadily increased, with the most stable G4s appearing closest to insertion breakpoints (Q21–Q35) **(Figure 5C)**. Comparable stability profiles for both IR and G4 motifs were also observed at deletion breakpoints, indicating that these breakpoint-associated patterns are not specific to insertion events (**S6 Fig A-B**).

**Figure 5.**
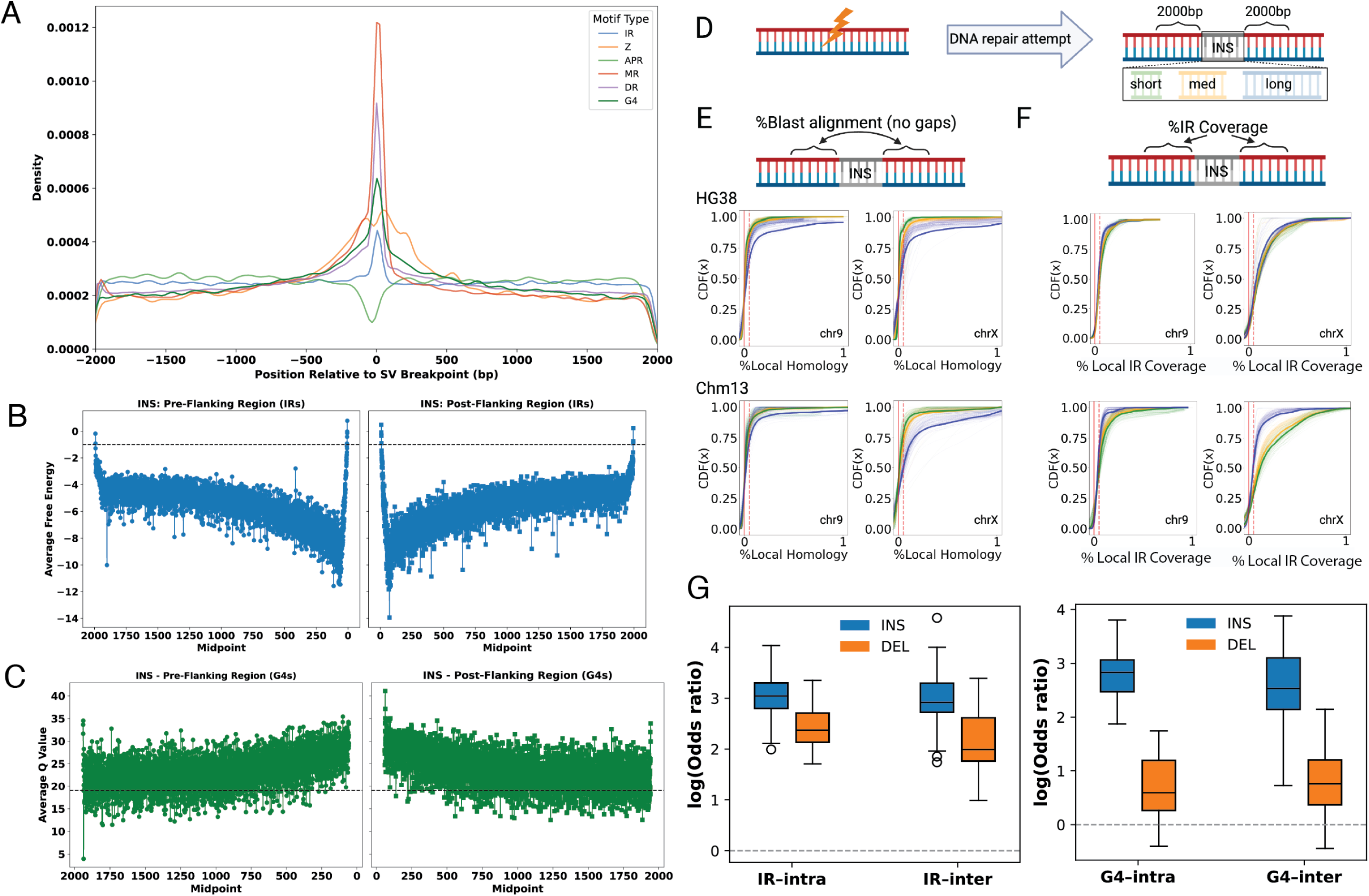
Non-B DNA motifs and structural variant (SV) flanking-region architecture. **(A)** Kernel density estimates (KDE) of six non-B DNA motif classes across ± 2kb surrounding insertion breakpoints. Multiple motif types show pronounced clustering at breakpoint positions, with the strongest peaks observed for mirror repeats (MRs) and direct repeats (DRs). **(B)** Average free energy profiles of inverted repeats in the 2 kb pre- and post-flanking regions of insertions, showing increasing IR stability as the breakpoint is approached, followed by a sharp local destabilization at the breakpoint. **(C)** Average stability (Quadron Q-scores) of G-quadruplex (G4) motifs across insertion flanking regions, demonstrating progressively higher G4 stability near breakpoints. **(D)** Schematic overview of the breakpoint-centered flanking-sequence analysis used to quantify local homology and non-B motif content across insertion classes. **(E)** Cumulative distribution functions (CDFs) of local BLAST alignment (left) and inverted repeat coverage (right) in insertion flanks from GRCh38 and T2T-CHM13 assemblies, stratified by insertion length (short, medium, long). **(F)** log odds ratios (Fisher’s exact test) for G4 and IR enrichment within insertions and deletions overlapping fixed intrachromosomal and inter-chromosomal segmental duplications (SDs).

The formation of SVs is often associated with underlying double-strand breaks (DSBs), which can result from various repair strategies that may be effective or ineffective. These repair mechanisms are strongly influenced by the extent of sequence homology around the SV breakpoint junction, with non-allelic homologous recombination (NAHR) requiring substantial lengths and quality of homology, whereas non-homologous end joining (NHEJ) requires little or no homology (53). Therefore, we extracted 2 kb flanking regions upstream and downstream of SV breakpoints to analyze local nBDNA patterns in regions near SVs (**Figure 5D**). We stratified insertions by their length—short, medium, and long—and compared their sequence homology and IR coverage using both GRCh38 and CHM13 mapped assemblies. In GRCh38 mapped assemblies, we observed that longer insertions on autosomes, such as chr9, tend to have lower local homology rates than shorter insertions (Kolmogorov-Smirnov (KS), p-value = 3.156e-08; Wasserstein distance (WS) = 0.043; **Figure 5E**). Conversely, IR coverage in the flanking regions of longer insertions was higher (KS p-value = 5.187e-16, WS = 0.018; **Figure 5F**). In CHM13-mapped assemblies, the homology pattern remained consistent (KS p-value = 0.0012, WS = 0.0169), although we observed increased divergence in IR coverage (KS p-value = 1.474e-13, WS = 0.0341). This divergence is more pronounced on sex chromosomes, such as chrX (GRCh38 WS = 0.0321 vs CHM13 WS = 0.1252). These results likely reflect the enhanced resolution of repetitive and structurally variable regions provided by the CHM13 assembly. Furthermore, since IRs may contribute to genomic fragility, repair outcomes—such as SV length—may be jointly influenced by both IRs and local homology.

Prior studies have also highlighted the enrichment of SVs within highly repetitive genomic regions containing SDs (54, 55). Therefore, we examined whether nBDNA motifs within SVs exhibit increased instability when overlapping with fixed SDs. We found that both G4 and IR motifs were highly enriched among SVs within fixed intra- and inter-chromosomal SDs compared to SVs outside these regions (**Figure 5G**). This enrichment was substantially greater for insertions than for deletions across all categories. For G4 motifs, enrichment within SD-overlapping SVs was strongly dependent on SV type (INS vs DEL). For example, G4s were enriched within insertions overlapping both intra-chromosomal (*x̃_logOR_intra_*: 2.83) and inter-chromosomal SDs (*x̃_logOR_inter_*: 2.53), whereas deletions showed substantially weaker enrichment (0.60 and 0.76, respectively). Similarly, IR motifs were highly enriched within insertions overlapping intrachromosomal SDs (*x̃_logOR_intra_*: 3.04) and inter-chromosomal SDs (*x̃_logOR_inter_*: 2.92), while deletions exhibited lower, though still elevated, enrichment (*x̃_logOR_intra_*: 2.38, *x̃_logOR_inter_*: 1.99). These findings indicate a synergistic instability effect, where inherently fragile non-B motifs are more concentrated in recombination-prone genomic regions. Not only are non-B DNA motifs more likely to be found within SVs, but they are also especially enriched in SVs located in SD regions, potentially acting as synergistic hotspots of genomic instability.

### Mobile Element Insertions Exhibit Family-Specific Enrichment and Stability Patterns of Non-B DNA Motifs

Beyond SVs, MEIs are another major source of genomic variation and instability. Therefore, we examined the relationship between nBDNA motifs and major mobile element families, including LINE/L1, SINE/Alu, and Retroposon/SVA elements. Here, motif coverage is defined as the total number of base pairs within a given insertion that overlap annotated non-B DNA motifs, averaged per mobile element family. Using this metric, we found that Retroposon/SVA insertions exhibited the highest nBDNA motif coverage on average across all motif types (*μ*_*SVA*_*coverage*_= 72bp), followed by LINE/L1 elements (*μ*_*L*1_*coverage*_= 43bp) and SINE/Alu elements (*μ*_*Alu*_*coverage*_= 30bp; **Figure 6A**). Notably, despite being shorter than LINE/L1 elements, SVAs still harbour a greater absolute number of nBDNA motif base pairs per insertion, indicating that this enrichment is not solely driven by element length. Within these SVA insertions, we found that DR (*μ*_*DR*_*coverage*_= 283bp), IR (*μ*_*IR*_*coverage*_= 255bp), and MR (*μ*_*MR*_*coverage*_= 203bp) motifs each occupy on average 200-280bp per element, while G4 and Z-DNA motifs occupy approximately 25bp.

**Figure 6.**
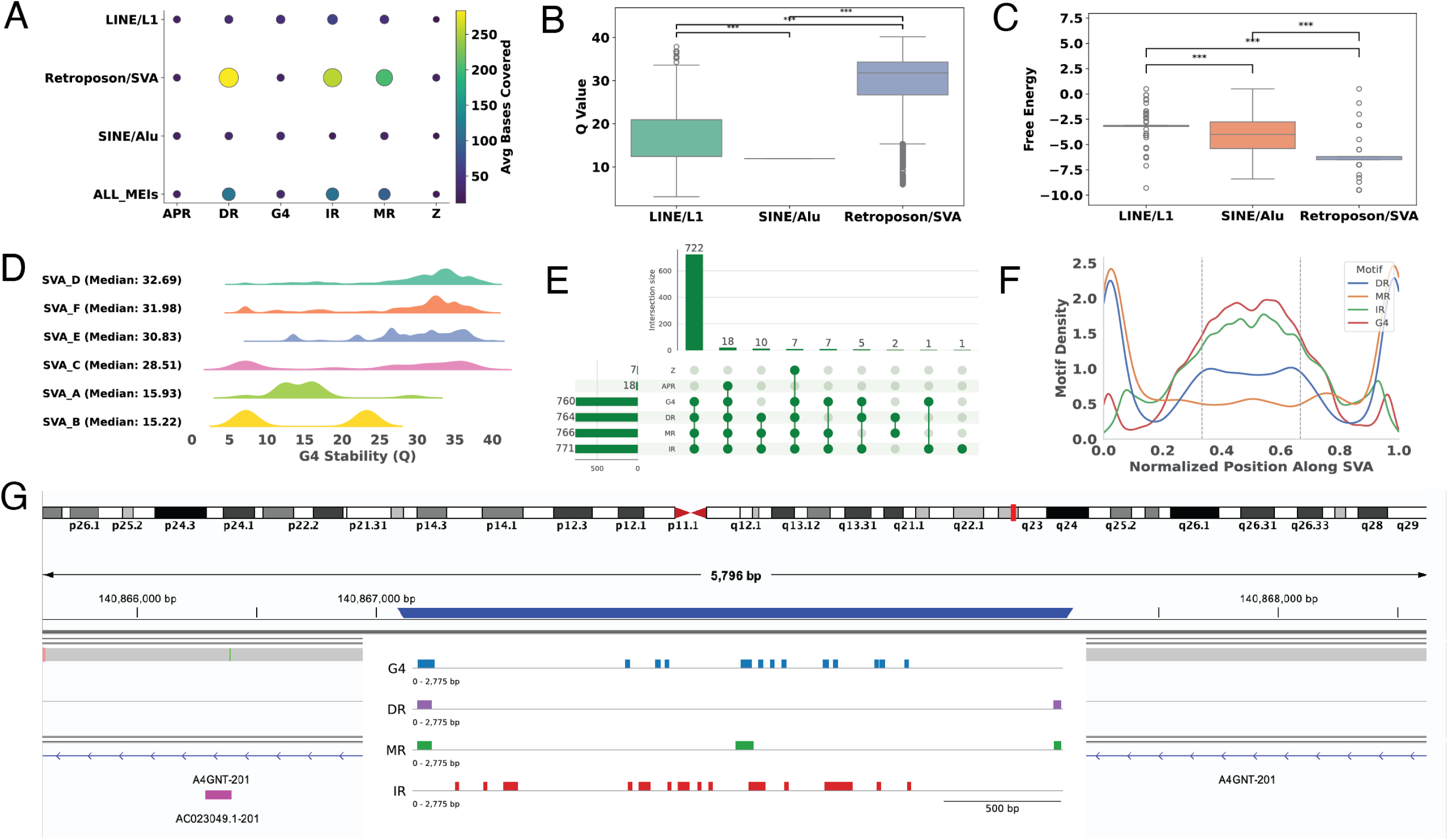
Family and sub-family specific non-B DNA motif composition, stability, and spatial organization within mobile element insertions (MEIs). (A) Average number of bases covered by each non-B DNA motif class within LINE/L1 SINE/AU, and Retroposon/SVA insertions. Retroposon/SVA elements display the highest overall motif coverage across all motif types. (B) Distribution of G4 stability (Quadron Q-score) across MEI families. Retroposon/SVA insertions harbour significantly more stable G4s than LINE/L1 and SINE/Alu elements (Mann-Whitney U, p < 0.001). (C) Free energy distributions of inverted repeats (IRs) across MEI classes, showing that SVA elements contain the most stable IR structures relative to other MEIs elements (Mann-Whitney U, p < 0.001). (D) Stability profiles of G4 motifs across SVA subfamilies (A-F), revealing that evolutionarily younger subfamilies (D-F) contain G4s with the highest predicted stability, whereas older families show lower and more variable stability. (E) UpSet plot showing recurrent combinations of co-occurring non-B DNA motifs within SVA elements. A total of 722 SVAs contain DR, MR, IR, and G4 motifs, the most common motif combination observed genome wide. (F) Normalized positional density of non-B DNA motifs along the canonical SVA structure. DR and MR motifs are enriched toward the 5’ and 3’ ends, while G4 and IR motifs are concentrated within the central FC-rich VNTR domain, with an additional small G4 peak in the CCCTCT-rich 5’ region. (G) Representative example of an SVA-F insertion at the A4GNT locus (present in 75 haplotypes), illustrating the dense clustering of motifs within an individual SVA.

These results align with the genetic architecture of SVA (SINE-VNTR-Alu) elements (56), specifically a 5’ CT-rich hexamer region, an antisense Alu-like sequence, a central GC-rich VNTR (variable number tandem repeat) domain with approximately 30-50 bp repeat units, a SINE-R component, and a 3’ poly-A tail (56). Therefore, we investigated whether the distinct architecture of SVA elements accounts for the enrichment of nBDNA motifs, compared to other MEI classes. In particular, the repetitive and GC-dense composition of these elements creates multiple sequence contexts that support various nBDNA-forming motifs. Consequently, we classified nBDNA motifs based on their stability profiles and found that Retroposon/SVA insertions contained the most stable G4 and IR structures compared to LINE/L1 and SINE/Alu elements (p-value < 0.001; **Figures 6B-C**). This pattern remained consistent within subfamilies, with the evolutionarily youngest SVA subfamilies (D, E & F) exhibiting G4s with the highest predicted stability scores. In contrast, older SVAs exhibited lower G4 stability and greater variability (Figure 6D). The distribution of G4s relative to age likely reflects selective pressures that maintain G4-forming potential in younger, more active elements, while older SVAs accumulate mutations that disrupt these structures (29). Furthermore, we leveraged CpG methylation annotation based on GRCh38 to provide a complementary epigenetic context (**S7 Fig. A**). We observed that G4 and IR motifs located within SVA insertions exhibit higher levels of CpG methylation compared to LINE/L1 and SINE/Alu elements. This suggests that, while SVAs harbor dense, intrinsically stable nBDNA motifs at the sequence level, their structural potential *in vivo* may be modulated by epigenetic repression.

Next, we examined the co-occurrence patterns of various nBDNA motifs within SVAs to determine whether certain nBDNA motif combinations are characteristic of specific SVAs. We identified 722 of 791 SVA insertions (91%) with overlapping DR, MR, IR, and G4 motifs, representing the most common motif combination across all SVAs **(Figure 6E)**. Spatially, among these SVAs, DR and MR motifs appear to cluster near the 5’ and 3’ ends, whereas G4s and IRs are enriched within the central VNTR region (**Figure 6F**). Additionally, we observed a small G4 peak near the 5’ end, consistent with previous reports indicating that G4s preferentially localize within both the central GC-rich VNTR and the CCCTCT-rich 5’ domain of SVA elements (29, 57). For example, one such SVA-F insertion in the A4GNT locus was present in 75 haplotypes in the CHM13-based HGSVC assemblies (**Figure 6G**), demonstrating the repetitive nature and high motif density of these SVA elements. Within the inserted element, we found nBDNA motifs reflecting the genome-wide density patterns observed across SVAs, such as the high density of DR and MR motifs in the flanks, G4 and IR motifs within the central VNTR domain, and G4s also present within the 5’ CCCTCT hexamer repeat at the 5’ end.

## Discussion

Recent advances in lrWGS and T2T assembly have enabled an unprecedented exploration of repetitive and structurally complex regions in the human genome. Leveraging these technological advances, large-scale initiatives such as HGSVC and HPRC now deliver comprehensive haplotype-resolved human genome assemblies from diverse populations, which were previously impossible to generate using srWGS techniques (6, 7). These resources enable systematic characterization of putative nBDNA sequence motifs across human populations and facilitate understanding of their contributions to genome organization, regulatory functions, and genomic instability. nBDNA structures are increasingly recognized as key regulators and drivers of human disease, with extensive evidence linking their formation to cancer-associated mutagenesis and genome rearrangements (24, 58, 59), to microlesions underlying inherited disorders (60), and even to emerging therapeutic targets in cancer treatment (61). In this study, we leveraged 130 phased haplotype-specific assemblies aligned to T2T-CHM13 and GRCh38 to generate a comprehensive population-scale map of six major non-B DNA motif classes. Our work provides a complete portrait of the non-B DNA landscape across human genomes assembled at near-complete resolution.

We first examined the genome-wide distribution of nBDNA motifs and found that although their cumulative coverage varied, their relative proportions were highly consistent across different haplotypes and populations. For instance, haplotypes aligned to CHM13 consistently exhibited greater cumulative motif content than their GRCh38-aligned counterparts, reflecting better representation of repetitive regions and the inclusion of nearly 200 Mb of previously unresolved sequence in T2T-CHM13 (1). However, this is likely an underestimate because our reference-based alignment approach excludes contigs or local haplotype configurations that fail to map to the reference, thereby excluding some population-specific or structurally divergent nBDNA sequences. Population-level comparisons showed that several motif classes, including IRs and MRs, had higher coverage in African haplotypes. This aligns with broader patterns of increased genetic variation observed in individuals from African populations (62, 63). Since IRs can form cruciform structures and MRs can cause slipped-strand intermediates, both of which promote replication fork stalling, slippage, and SV formation, their increased coverage likely reflects the greater population diversity and genomic variation found in African genomes. Meanwhile, G4 densities remained relatively stable across populations, possibly indicating that certain nBDNA motifs are subject to stronger evolutionary or functional constraints than others.

Next, we performed systematic biophysical stability predictions to understand how the sequence stability of nBDNA motifs relates to their genomic distribution. Our regression analyses showed that repeat length (arm length) was a stronger determinant of stability, with spacer length exerting a much smaller effect. This is consistent with established biophysical models in which cruciform extrusion becomes energetically favorable when IRs contain long arms (40, 64). In contrast, G4 stability was primarily influenced by GC content and loop length architecture. This pattern is expected because higher GC content promotes strong G-quartet stacking, and shorter loops promote more compact, parallel G4 topologies that are inherently more stable than G4s with longer loops; however, the strength of this relationship varies with the presence of specific cations (65, 66).

When combined with regulatory annotations, nBDNA motif enrichment exhibited highly context-dependent patterns. For example, IRs were not widely enriched across cis-regulatory elements, consistent with previous reports (16). However, when stratified by predicted stability, we found that stable cruciform-forming IRs were selectively retained within a small subset of promoter-like signatures, including notable loci such as the CDKN1A promoter. This suggests that while IRs are not globally enriched in promoters, high-stability IRs may be conserved at functionally important sites where cruciform extrusion plays a regulatory or structural role. This aligns with prior evidence that cruciforms are recognized by proteins involved in transcription and genome stability (e.g., p53, BRCA1, PARP-1, RAD54, MLL, WRN) (44). In contrast, G4 motifs showed strong enrichment across cCREs, particularly in promoter-like (PLS) and proximal enhancer-like (pELS) elements. Stratifying by predicted stability revealed that unstable G4s were more enriched in CTCF-only and distal enhancer (dELS) regions, consistent with the instability-biased G4 profiles observed in a prior study (67). These patterns are biologically plausible since stable G4s tend to localize to highly active, GC-rich regions immediately flanking transcription start sites (16, 67). The strong preference of stable G4s for promoter-proximal, transcriptionally active DNA may explain the minimal enrichment observed in CTCF-only and dELS regulatory elements, both of which are usually located further from the TSS than other elements.

Similarly, fully assembled centromeres in our dataset were densely populated with multiple classes of nBDNA motifs consistent with a previous study (68). However, as observed in great apes (36), no single motif class showed consistent or universal enrichment across chromosomes or haplotypes. Instead, the nBDNA landscape was highly variable, likely reflecting differences in centromere sequence composition and HOR organization. One exception was the G4 motif, which was consistently depleted across all centromeres. A probable explanation is that human alpha-satellite DNA is strongly AT-rich (69, 70), creating sequence contexts that support inverted, direct, and mirror repeats but contain few G-rich tracts capable of forming stable G4s. Beyond global centromeric patterns, our analyses revealed localized enrichment of specific nBDNA motif classes within CENP-B boxes on certain chromosomes. Existing models suggest that nBDNA and CENP-B binding motifs can functionally replace each other at centromeres (68). In contrast to this purely compensatory view, our findings showed chromosome-specific cases in which specific nBDNA motifs were enriched within and around CENP-B boxes. Although this does not imply direct causation, these observations argue against a strictly compensatory model in which nBDNA structures and CENP-B binding are mutually exclusive, instead suggesting that both features can co-occur within specific centromeric contexts.

Expectedly, highly repetitive segmental duplications also showed clear and motif-specific enrichment patterns in our study, with DRs, G4s, and APRs consistently enriched across both intra- and inter-chromosomal fixed SDs. Since we focused only on invariant SDs (duplications fixed across all human haplotypes), the enrichment we observe likely reflects stable and long-lasting duplication structures rather than the more dynamic variable SDs that occur in specific individuals. Previous studies have demonstrated that inter-chromosomal SDs are often invariant and frequently located in sub-telomeric and pericentromeric regions (51), which are known to contain tandemly repeated satellites that support the formation of multiple nBDNA structures. In contrast, we observed little enrichment of MRs and IRs within all fixed SDs, even though both motif classes are known to form highly unstable cruciform and triplex H-DNA, which can promote breakage and large-scale chromosomal rearrangements (71, 72, 73, 74). Earlier, it was reported that rare SDs tend to be longer and show unusually high sequence identity (51). Since MRs and IRs are known to contribute to large-scale chromosomal rearrangements, the absence of enrichment for these motifs in our results likely reflects our focus on fixed rather than rare SDs, which prevents us from capturing motifs that may specifically contribute to the formation of these longer, rarer duplication events. Unlike IRs and MRs, G4s showed consistent enrichment within fixed SDs and were elevated within intrachromosomal duplication blocks. This enrichment aligns with previous findings that GC-rich repeat elements are directly involved in the formation and pericentromeric localization of both inter- and intra-chromosomal SD events (75). The enrichment of DR, G4, and APR motifs, alongside the lack of IR and MR enrichment, suggests that only certain nBDNA structures are tolerated within evolutionarily older SDs, while more destabilizing motifs may be confined to rare or more recently evolved duplication events that our analysis of the fixed SD set did not capture.

Enrichment analysis of nBDNA motifs around SV breakpoints provided further insights into their role in promoting genomic variations. Across haplotypes, we observed distinct clustering of nBDNA motifs at SV boundaries, with DRs and MRs showing the strongest enrichment, and stable G4 motifs accumulating immediately at the breakpoints. In contrast, IRs became increasingly stable as they approached the breakpoint, yet showed a marked loss of stability exactly at the SV junction in both insertions and deletions. These opposing patterns support the idea that different repair pathways act on each motif type. IR-induced cruciforms can generate double-strand breaks, which are often resolved by microhomology-mediated end joining, an error-prone pathway that resects DNA and disrupts the original inverted symmetry at the repaired junctions (76, 77). Conversely, G4-induced replication stalls are typically processed via homologous recombination, a template-guided mechanism that preserves the GC-rich sequence and can maintain highly stable G4s at breakpoints (78). The contrasting stability patterns we observe—IRs becoming unstable at the junction while G4s remain highly stable—may reflect the distinct processing outcomes of MMEJ and HR during SV formation. Notably, longer insertions showed greater IR coverage in their flanking regions without a corresponding increase in local homology, consistent with MMEJ-driven repair, since IR-induced hairpins can produce double-strand breaks repaired by microhomology rather than homology-driven recombination. Finally, SVs within fixed SDs showed strong enrichment for both G4s and IRs compared to SVs outside these duplicated regions. Because SDs continue to accumulate substitutions, indels, and retrotransposition events long after their formation (54, 55), SVs arising within these regions occur against ongoing sequence turnover, predisposing them to instability. The presence of nBDNA motifs within these duplicated regions likely amplifies, rather than independently causes, susceptibility to breakage and aberrant repair in these areas.

Similarly, MEIs, particularly SVAs, exhibited the highest densities of nBDNA motifs (including DRs, IRs, MRs, and G4 motifs), often co-occurring within the same element. This is consistent with the well-documented architecture of SINE-VNTR-ALUs (SVAs), whose GC-rich VNTR domains and repetitive 5’ CCCTCT hexamer region create abundant sequence contexts permissive for multiple nBDNA structures (29, 79). Our finding that evolutionarily younger SVA subfamilies (SVA D-F) harbor the most stable G4s aligns with previous studies showing that nBDNA, especially G4s, may play active roles in the MEI life cycle by affecting transcriptional activation, retrotransposition efficiency, and R-loop formation during reverse transcription (29, 30). nBDNA motifs have also been linked to shaping MEI integration preferences, and the high density and stability of G4s and IRs within SVAs may contribute to their previously observed insertional biases and association with genomic instability hotspots (80, 81). More broadly, our observation that SVAs contain the highest abundance of overlapping DR, MR, IR, and G4 motifs supports the idea that MEIs act as key mechanisms for spreading nBDNA across the genome, thereby affecting local chromatin environments, modulating promoter activity, and possibly facilitating heterochromatin formation around young insertions (9, 35, 82, 83). Finally, the spatial clustering we observe—where DRs and MRs are more concentrated at SVA boundaries and G4 and IR motifs are enriched within the GC-rich VNTR core—mirrors the functional division proposed for nBDNA structures in earlier MEI studies (80,81). Boundary-localized DR and MR motifs may enhance recombination and integration-related instability, while VNTR-localized G4s and IRs are positioned to influence retrotransposition efficiency, reverse transcription dynamics, and regulatory interactions.

Taken together, our study demonstrates that nBDNA motifs are intricately woven into the structural and regulatory architecture of human genomes. Their distribution across haplotypes is affected by sequence composition, population-level differences, evolutionary history, and local genomic context. Their enrichment for regulatory elements underscores their potential to influence transcriptional activity, whereas their concentration at centromeres, SDs, SV breakpoints, and MEIs highlights their roles in maintaining structural integrity and promoting genetic variation. Although our study provides a comprehensive overview of non-B DNA motifs across high-quality human genomes, several limitations should be acknowledged. Our analyses rely on a reference-based approach, in which high-quality haplotype assemblies are aligned to reference genomes to ensure consistent annotation across individuals. This method restricts our ability to detect highly divergent or haplotype-specific sequences that do not align well with the reference genome. Future research using fully de novo haplotype assemblies and their associated annotations will be essential to fully characterize the population-specific landscape of nBDNA motifs. Additionally, motif annotations were generated using sequence-based prediction frameworks that identify regions capable of forming secondary structures; however, we cannot confirm their in vivo folding. Although integrating biophysical stability predictions improves these estimates, experimental validation remains necessary to verify which motifs fold under physiological conditions. Furthermore, many SVAs in the human genome are epigenetically silenced by methylation (84, 85), and methylation directly suppresses G4 formation and stability (86). Although we incorporated Nanopolish-derived methylation analyses, these data remain reference-based and are not completely resolved at the haplotype level. Future integration of haplotype-resolved methylation profiles will be essential to further explore how epigenetic differences among individuals influence nBDNA formation, stability, and activity within MEIs. Moreover, our classification of MRs did not differentiate among higher triplex-forming potentials, and we did not include short tandem repeats (STRs) as an additional category of potential nBDNA motifs, which are important sources of structural diversity and warrant exploration in future studies. Lastly, our interpretations of regulatory enrichment are based on ENCODE cCRE annotations derived from GRCh38, which reflect averaged, cell-type-agnostic regulatory landscapes rather than tissue-specific patterns. Consequently, subtle, context-dependent variations in nBDNA motif enrichment across developmental stages, tissues, or cellular states are not accounted for in our study.

Despite these limitations, our study develops a comprehensive haplotype-resolved atlas of nBDNA motifs across diverse human genomes. By integrating motif prediction, stability modeling, regulatory annotation, and structural variation mapping across T2T assemblies, our work offers a framework for understanding how non-canonical secondary structures influence genome organization, regulatory complexity, and genomic instability. As long-read sequencing becomes more accessible and de novo T2T assemblies expand to include larger and more diverse populations, these findings will help guide efforts to interpret repetitive DNA and uncover the molecular mechanisms by which nBDNA shapes human genetic diversity.

## Materials & Methods

We utilized 130 high-quality haplotype-resolved human assemblies from the HGSVC3 Project (7). These samples were sequenced using PacBio Hi-Fi and ultra-long Nanopore long-read technology, assembled with the Verkko algorithm, and aligned to both the CHM13v2.0 and GRCh38.p14 reference genomes. HGSVC3 serves as a resource of telomere-to-telomere, high-quality, diverse human assemblies, aligned to both reference genomes using minimap2 v2.26 (7).

### Annotating Non-B DNA Motifs Across Long-Read Human Assemblies Aligned to CHM13v2.1 and GRCh38.p14 References

We annotated the 130 phased haplotype assemblies that were aligned to both T2T-CHM13v2.0 and GRCh38.p14 reference genomes. For each haplotype, we selected primary alignments (samtools -F 256) to retain only well-mapped, high-quality reads. This filtering ensured that motif annotation was based on the most reliable alignments, which is essential given the repetitive nature of nBDNA motifs. After partitioning the BAM file by chromosome with bamtools split, we converted the resulting files to FASTA format with samtools fasta for computational efficiency. Subsequently, chromosome-level FASTA files were merged per haplotype to generate complete haplotype assemblies, which were annotated with both nBMST (31) and Quadron (38) to identify non-B DNA motifs. nBMST identifies canonical non-B motifs based on defined sequence patterns and was used to annotate IRs, DRs, MRs, APRs, and Z-DNA motifs using default parameters. Quadron applies a machine-learning framework (a tree-based gradient-boosting model) trained on experimental G4-seq data to predict G4 structures and an associated stability score based on polymerase stalling (Q-score). We used a stability score of 19 to denote stable G4s (38). However, we considered all predicted G4s, enabling stability-based analyses. The resulting annotations from both tools were collapsed into non-redundant motif sets using bedtools merge.

### Quantifying Structural Stability and Functional Context of G-Quadruplexes and Inverted Repeats

We utilized known regulatory sites to investigate the presence, location, and stability of the nBDNA motifs, IR, and G4. For G4 motifs, we utilized the Quadron-derived stability scores (Q-score), whereas for IR motifs, we quantified using the minimum free energy (MFE) predicted by seqfold. The seqfold tool implements the Zuker (1981) (87) dynamic-programming algorithm, in which lower (more negative) free-energy values correspond to greater thermodynamic stability. We classified IRs with MFE < 0 kcal/mol as stable and those with MFE > 0 kcal/mol as unstable. While repeat and spacer thresholds are typically used to define IR stability, we prioritized a biophysical definition based on predicted folding energetics, thereby moving beyond criteria for repeat and spacer lengths.

We obtained candidate cis-regulatory element (cCRE) annotations from the ENCODE Project (v3), which are defined on the GRCh38 reference genome and available through the SCREEN Encode portal (43). This is a unified cCRE dataset that integrates DNAase-seq accessibility, histone modification, and CTCF-binding data from experiments across diverse cell and tissue types and classifies regulatory elements into promoter-like (PLS), proximal enhancer-like (pELS), distal enhancer-like (dELS), CTCF-only (CTCF), and DNAse-H3K4me3 cCREs (43). To evaluate the regulatory distribution of nBDNA motifs, we intersected motifs with ENCODE cCREs using bedtools intersect -*wo* to count the number of base pairs a nBDNA motif sequence overlaps with each cCRE class. We first quantified genome-wide enrichment of nBDNA motifs within cCREs without stratification by stability. For each motif type and cCRE category, we constructed 2×2 contingency tables comparing nBDNA motif bases overlapping cCREs versus non-overlapping motif base pairs across the genome, and calculated odds ratios using Fisher’s exact test.

To evaluate how structural stability influences regulatory localization, we categorized motifs into stable and unstable groups and conducted within-haplotype comparisons of these motifs overlapping each cCRE class. For each haplotype, motif type, and cCRE category, Fisher’s exact test was applied to calculate odds ratios that measure the relative enrichment of stable versus unstable motifs. These odds ratios were log-transformed for further analysis and visualization. Moreover, we calculated p-values and 95% confidence intervals for haplotype-level odds ratios for enrichment analyses (**Supplementary Tables 4-10**).

### Annotation of Non-B DNA Motifs Within Centromeric and Alpha-Satellite Higher-Order Repeat Arrays

To investigate the organization and structural context of nBDNA motifs within human centromeres, we annotated centromeric and alpha-satellite regions across the haplotype assemblies. These assemblies include complete, base-level-resolved centromeres that are fully assembled through alpha-satellite higher-order repeat (HOR) arrays, representing some of the most repetitive regions of the genome (7). Importantly, any contigs that spanned the centromere or contained misassemblies (misjoins, collapses, and false duplications) were excluded, ensuring that only high-confidence, correctly assembled centromeres were analyzed (7). We obtained the bed files from the HGSVC3 resource. We converted the provided coordinates into haplotype-specific bed files and extracted the corresponding centromeric sequences from the unaligned haplotype FASTA assemblies using seqtk. The extracted centromeric and HOR sequences were annotated with nBDNA motifs using nBMST and Quadron (Q > 19), and overlapping motif coordinates were collapsed using bedtools merge to remove redundant nBDNA motifs. All annotation outputs were processed to compile nBDNA motif information across complete centromeric regions and active alpha satellite higher-order repeat arrays for all haplotypes.

To further resolve centromeric architecture, we applied the Genomic Centromere Profiling (GCP) workflow (46). We utilized the *fuzznuc* module to identify CENP-B box motifs within each centromere. A CENP-B box is defined by a 17bp sequence (5’-NTTCGNNNNANNCGGGN-3’) which can be specifically bound by the CENP-B protein (46). This protein is the only centromeric protein known to recognize and bind a defined DNA sequence within alpha-satellite higher-order repeat (HOR) arrays (46, 88, 89). We intersected our nBDNA motifs in centromeres with the CENP-B box coordinates using `bedtools intersect -wo` to quantify the number of nBDNA motif bases overlapping CENP-B box motifs within each haplotype. We assessed enrichment using a Fisher’s exact test to compare base counts of overlapping nBDNA motifs within CENP-B boxes, to those outside CENP-B boxes.

We subsequently used the *model1* module of the GCP workflow to assign ∼171 bp alpha satellite monomeric units, and then partitioned HOR arrays into their constituent monomers to enable monomer counting. We intersected the nBDNA motif annotations with these HOR partitions again, using `bedtools intersect -wo` to calculate motif-based overlap per monomer class. Finally, we performed Fisher’s exact tests to determine whether specific nBDNA motifs were significantly enriched within particular monomeric contexts relative to the rest of the HOR array.

### Quantifying Non-B DNA Motif Enrichment and Composition in Segmentally Duplicated Regions

To examine the distribution of nBDNA motifs within segmentally duplicated genomic regions, we analyzed segmental duplications (SDs) that were fixed (i.e., common) across all 170 haplotype-resolved assemblies (including the CHM13v2.0 reference genome) from HGSVC and HPRC. This fixed SD callset was derived from 53 HGSVC and 47 HPRC samples, each sequenced and assembled with hifiasm, and segmental duplications were identified using SEDEF (51). To preserve haplotype specificity while ensuring consistency in SD annotation, we restricted our analyses to the subset of 53 HGSVC haplotypes for which CHM13-projected SD coordinates were available. Since nBDNA motif annotations were generated from CHM13-based alignments, this ensured one-to-one correspondence between each haplotype’s specific motif landscape and the fixed SD intervals. For consistency, we restricted our analyses to autosomes, given ploidy differences and assembly limitations on the Y chromosome, and excluded the short arms of acrocentric chromosomes (13, 14, 15, 21, and 22), where accurate assembly and mapping of SDs remain unreliable.

We analyzed fixed SDs categorized as intra- and inter-chromosomal, following the definitions established in a previous study (51). We denoted intra-chromosomal SDs as duplicated sequences within the same chromosome, whereas inter-chromosomal SDs are duplications spanning different chromosomes. Across the human genome, this fixed SD call set comprised 1,781 intra-chromosomal and 2,541 inter-chromosomal SDs, representing the most evolutionarily stable duplication regions shared among diverse human haplotypes. Fixed SDs were identified by merging per-haplotype SD annotations for each category (intra- and inter-chromosomal) and by calculating per-base coverage across all haplotypes in the CHM13 coordinate space using BEDTools and GenomeCov. Genomic intervals with full coverage across all assemblies (depth equal to the total number of haplotypes) were retained as “fixed,” indicating duplications being present in every analyzed haplotype and the CHM13 reference. Intra- and inter-chromosomal SDs were analyzed separately due to their distinct genomic configurations and evolutionary origins. nBDNA motif annotations from CHM13-aligned haplotype assemblies were intersected with SDs using bedtools intersect to determine the number of motif base pairs within versus outside fixed SDs. Genome-wide enrichment of each motif class within SDs was assessed for each haplotype using Fisher’s exact test by comparing the proportion of motif base pairs overlapping SDs to their distribution across the remainder of the genome.

To evaluate the effect of structural stability on SD localization, G4 and IR motifs were further divided into stable and unstable groups. Comparisons between these groups were performed within each haplotype and SD category. For each haplotype, Fisher’s exact test was used to determine odds ratios that measured the relative enrichment of stable motifs within SDs. These odds ratios were log-transformed for further analyses and visualization. Additionally, we calculated p-values and 95% confidence intervals for all haplotype-level enrichment statistics (**Supplementary Tables 1-3).**

### Characterizing Non-B DNA Motif Enrichment Within and Flanking Structural Variants (SVs) and Mobile Element Insertions (MEIs)

We investigated the relationship between non-B DNA motifs and genome instability by examining their enrichment proximal to SV breakpoints and MEIs across the haplotype-resolved human assemblies from the HGSVC. Haplotype SVs were generated using the Phased Assembly Variant Caller (PAV 2.4.0.1) as part of the HGSVC3 study (62). The callset of SVs includes phasing, denoting the SV genotype, whereby the first and second alleles correspond to the first and second haplotypes, respectively. Using this phased information, we separated the SVs into haplotype-specific BED files, yielding distinct variant sets for each haplotype analyzed. In contrast, the MEI call set was generated using a complementary MEI pipeline, which includes L1ME-AID, PALMER2, and MELT-LRA (7, 90). These callers jointly identified Alu, LINE-1, and SVA insertions and provided phased genotype information, which enabled us to split the MEI callset into haplotype-specific BED files.

For each haplotype, we created 2kb flanking intervals upstream and downstream of each SV and MEI breakpoint(s) to obtain sufficient sequence context surrounding the junction. This sequence extraction was done directly from the CHM13-aligned primary BAM files using samtools consensus -r “$region” –show-ins no -a “$bam_file” –mode simple. This process generated haplotype-specific flanks for each event, excluding inserted bases, while preserving the surrounding genomic context. To determine sequence homology between these flanks and gauge potential repair strategies, we performed gapless BLAST alignments between each pair of upstream and downstream flanks. Alignments were generated using a word size of 5 and required opposing orientations. Homologous sequences retrieved from BLAST were converted to FASTA files, then subsequently used for nBDNA motif annotation via nonB-GFA and Quadron. Since MEI flanks showed little enrichment for nBDNA motifs, we also annotated within-event sequences, including the inserted and deleted segments of the SV itself, to assess motif composition and stability.

To examine segmental duplication (SD) architecture, we used the fixed intra- and inter-chromosomal SD annotations described above, then identified the intersection between haplotype-specific SVs and nBDNA motifs. We compared the frequencies of nBDNA motifs between SVs that overlapped SDs and those that did not, to assess whether motifs are preferentially enriched in duplicated regions. This comparison allowed us to evaluate whether the coexistence of two instability-prone features (nBDNA motifs and SDs) marks regions of potential compounded fragility.

## Source code & Data Availability

The phased haplotype assemblies generated by the Human Genomic Structural Variation Consortium (HGSVC3), along with the associated centromere, SV, and MEI calls were obtained from the IGSR data portal (https://ftp.1000genomes.ebi.ac.uk/vol1/ftp/data_collections/HGSVC3/working/ & https://ftp.1000genomes.ebi.ac.uk/vol1/ftp/data_collections/HGSVC3/release/Mobile_Elements/1.0/ & https://ftp.1000genomes.ebi.ac.uk/vol1/ftp/data_collections/HGSVC3/release/Variant_Calls/1.0/ & https://ftp.1000genomes.ebi.ac.uk/vol1/ftp/data_collections/HGSVC3/release/Centromeres/1.0/). We downloaded alignments for Verkko assembled haplotypes only (PRJEB76276).

ENCODE cCRE annotations were accessed through the SCREEN (v3) portal (https://screen.encodeproject.org/)

Non-B DNA motif annotation tools used in this study are publicly available. Quadron for G4 prediction is available at (https://github.com/aleksahak/Quadron), Non-B Motif Search tool (nBMST) is available at (https://github.com/abcsFrederick/non-B_gfa). Seqfold for thermodynamic predictions of nucleic acid secondary structure free energy is available at (https://github.com/Lattice-Automation/seqfold). The Genomic Centromere Profiling (GCP) pipeline used for CENP-B annotation and HOR organization profiling is available at (https://github.com/GiuntaLab/GCP-Centeny/tree/main)

All relevant source codes for this study, including scripts for annotation and downstream analyses, can be accessed on the project GitHub page (https://github.com/kumarlab-compomics/PhasedHapAssembly-nonB)

## Acknowledgements

This research was made possible by the availability of high-quality haplotype-resolved genome assemblies generated by the Human Genomic Structural Variation Consortium (HGSVC). We thank Dr. Kateryna Makova for valuable discussions and expert insights on non-B DNA structures and genome evolution, which helped shape some of the interpretations presented in this work.

