## Supplementary figures for "Diversity and Genomic Organization of Non-B DNA Motifs in Haplotype-Resolved Human Genome Assemblies"

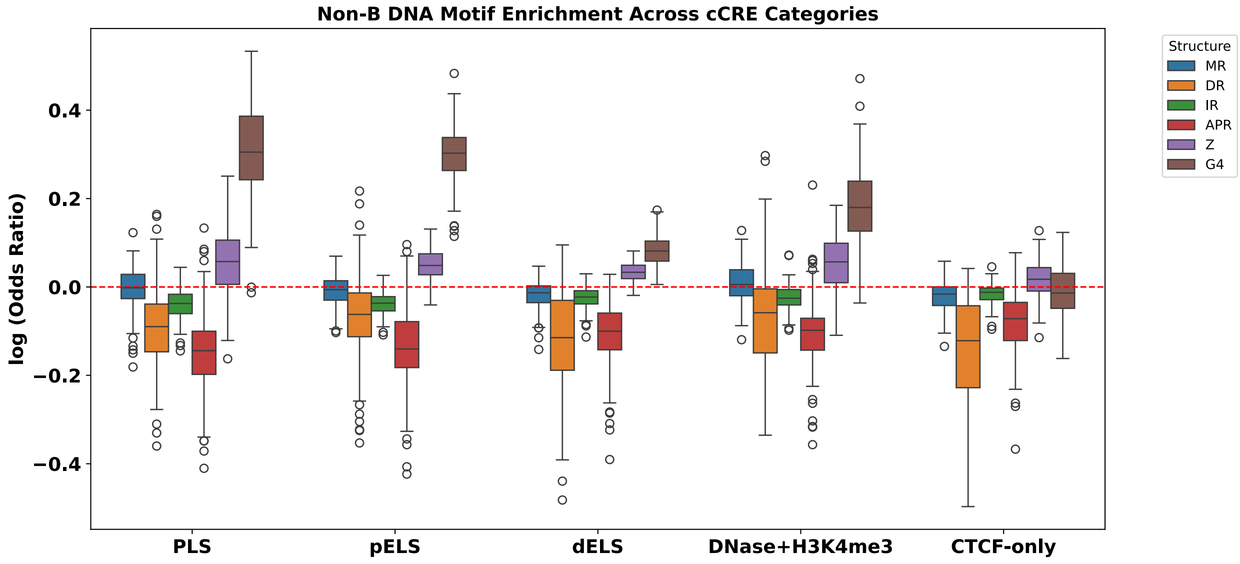


**Supplementary Figure S1. Non-B DNA motif enrichment across candidate cis-regulatory elements (cCREs).** Boxplots show haplotype-level log odds ratios for enrichment of six non-B DNA motif classes (MR, DR, IR, APR, Z-DNA, and G4) within ENCODE-defined cCRE categories including promoter-like (PLS), proximal enhancer-like (pELS), distal enhancer-like (dELS), DNase-H3K4me3, and CTCF-only elements. For each haplotype, enrichment was calculated using Fisher's exact test by comparing motif overlap within cCREs to the whole genomic background. The dashed red line indicates a log odds ratio of 0 (no enrichment).

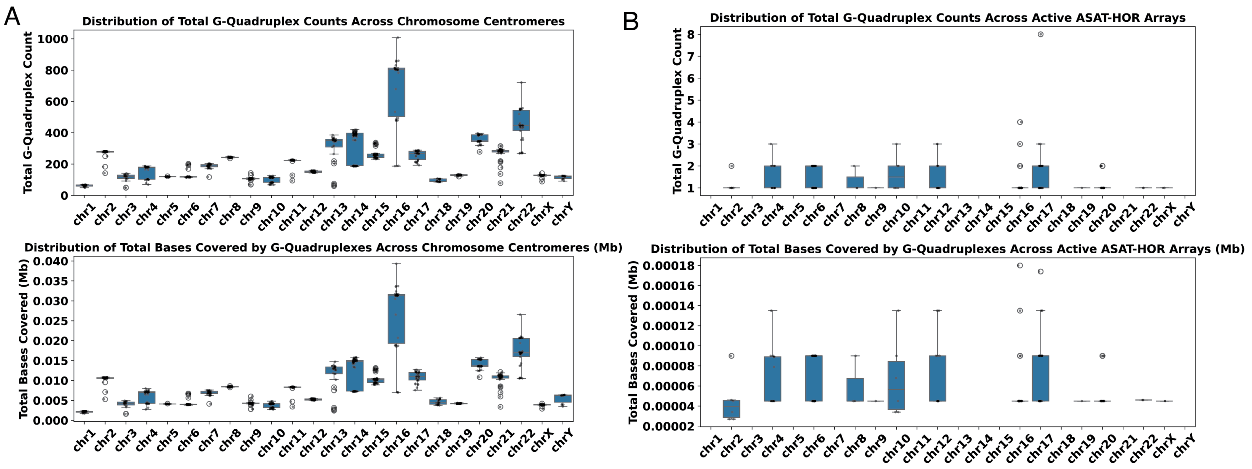


**Supplementary Figure S2 A-B. G4 abundance and base coverage within fully assembled human centromeres. (A)** Distribution of total G4 motif counts (top) and total base pairs covered by G4 motifs (bottom) across complete chromosome centromeric regions. **(B)** Distribution of total G4 motif counts (top) and total base pairs covered by G4 motifs (bottom) restricted to active α-satellite higher-order repeat (ASAT-HOR) arrays within centromeres. For each chromosome, boxplots summarize haplotype-level variation in G4 abundance across assemblies. Both complete centromeres and active ASAT-HOR arrays contain relatively few G4 motifs across all haplotype assemblies.

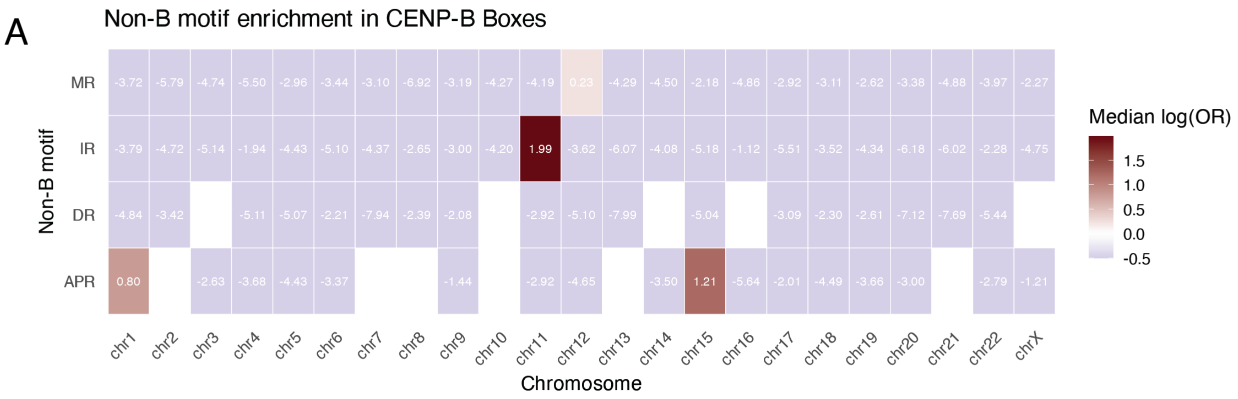


**Supplementary Figure S3. Chromosome-specific enrichment of non-B DNA motifs within CENP-B boxes.** Heatmap showing the median log odds ratios for enrichment of non-B DNA motif classes within CENP-B box sequences across individual chromosomes. For each haplotype, overlap between annotated non-B DNA motifs and CENP-B boxes was quantified at the base-pair level and enrichment was assessed using Fisher's exact test relative to centromeric background sequence outside CENP-B boxes. Values shown represent the median log odds ratio across haplotypes for each motif-chromosome combination. Cells with no values indicate motif-chromosome combinations for which no overlap between non-B DNA and CENP-B boxes was observed.

A

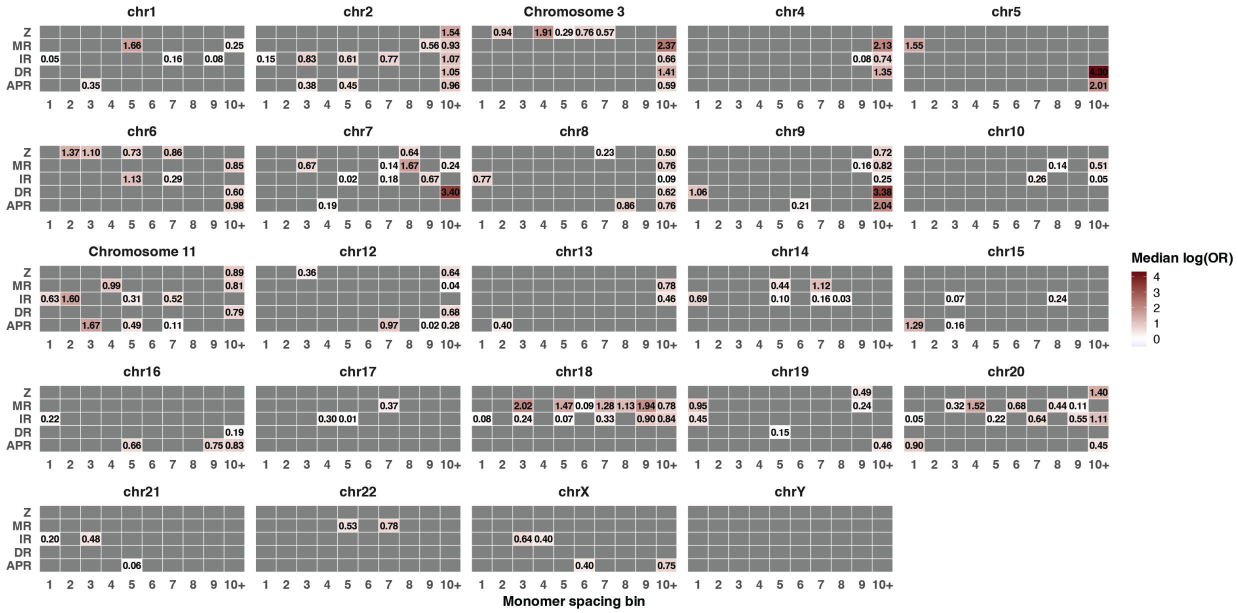

**Supplementary Figure S4. Non-B DNA motif enrichment in CENP-B monomer spacing intervals across human centromeres.** Heatmaps show the median log odds ratios for enrichment of non-B DNA motif classes within centromeric regions stratified by CENP-B box monomer spacing across individual chromosomes. CENP-B boxes were assigned to α-satellite higher-order repeat arrays and grouped by the number of monomer units separating consecutive CENP-boxes (monomer spacing bins 1-10+). For each haplotype and chromosome, enrichment of non-B DNA motifs within each spacing category was calculated using Fisher's exact test relative to centromeric background sequence. Values represent the median log odds ratio across haplotypes for each motif-spacing chromosome combination. Cells with no displayed value indicate spacing categories for which no overlap between non-B DNA motifs and the corresponding spacing bin was observed.

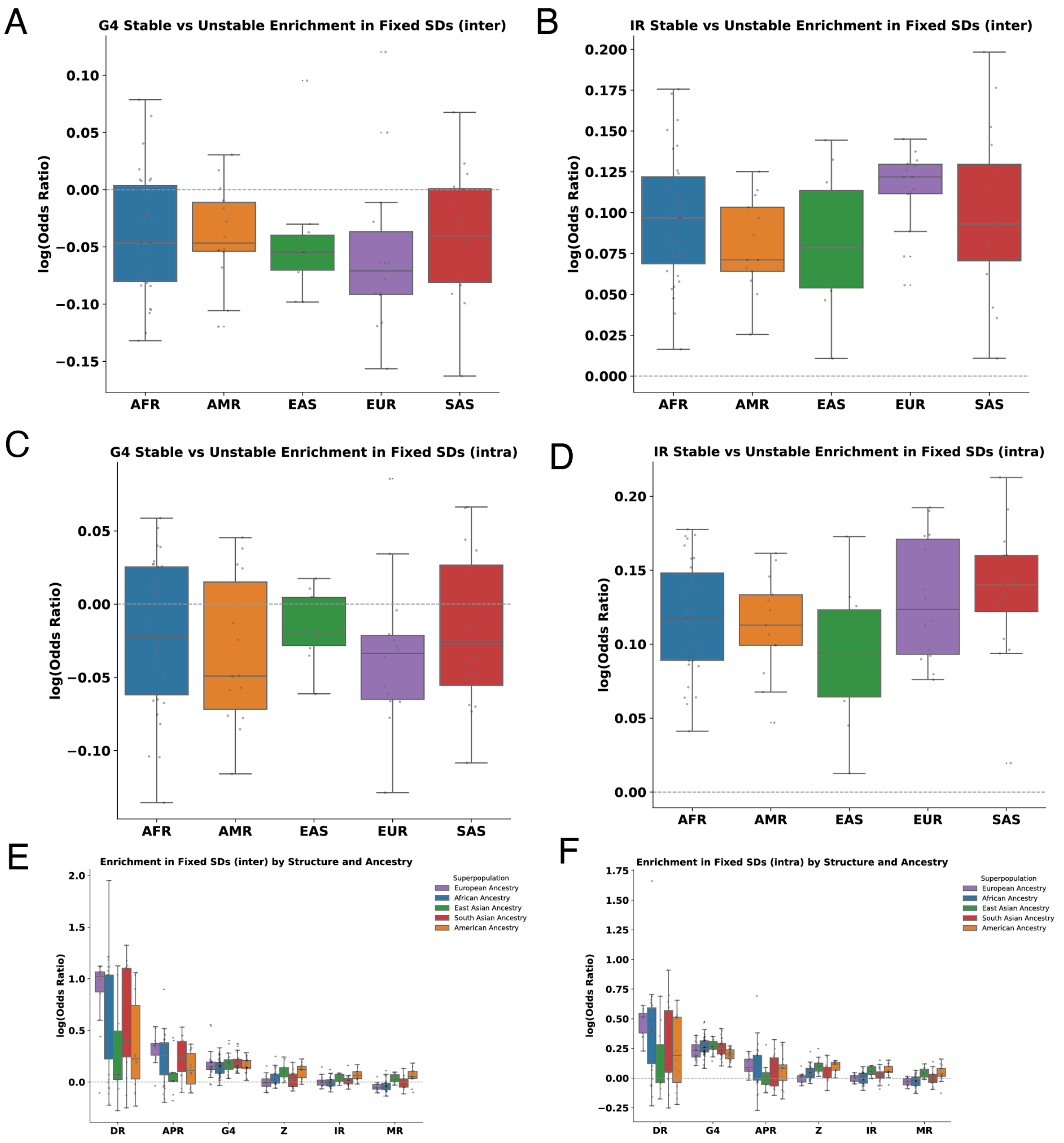


**Supplementary Figure S5 A-F. Population stratitifed enrichment of non-B DNA motifs within fixed segmental duplications. (A-B).** Boxplots show haplotype-level log odds ratios for enrichment of stable and unstable G4 motifs (A) and IR motifs (B) within fixed interchromosomal segmental duplications, stratified by population supergroup (African, East Asian, European, American, and South Asian). **(C-D).** Corresponding enrichment analyses for stable and unstable G4s (C) and IRs (D) within fixed intrachromosomal segmental duplications. For all panels A-D, enrichment was calculated per haplotype using Fisher's exact test by comparing motif overlap within fixed SDs to the genomic background. **(E-F).** Genome-wide enrichment of all non-B DNA motif classes within fixed interchromosomal (E) and intrachromosomal (F) segmental duplications, stratified by population supergroup. Across all panels, the dashed horizontal line indicates a log odds ratio of 0 (no enrichment).

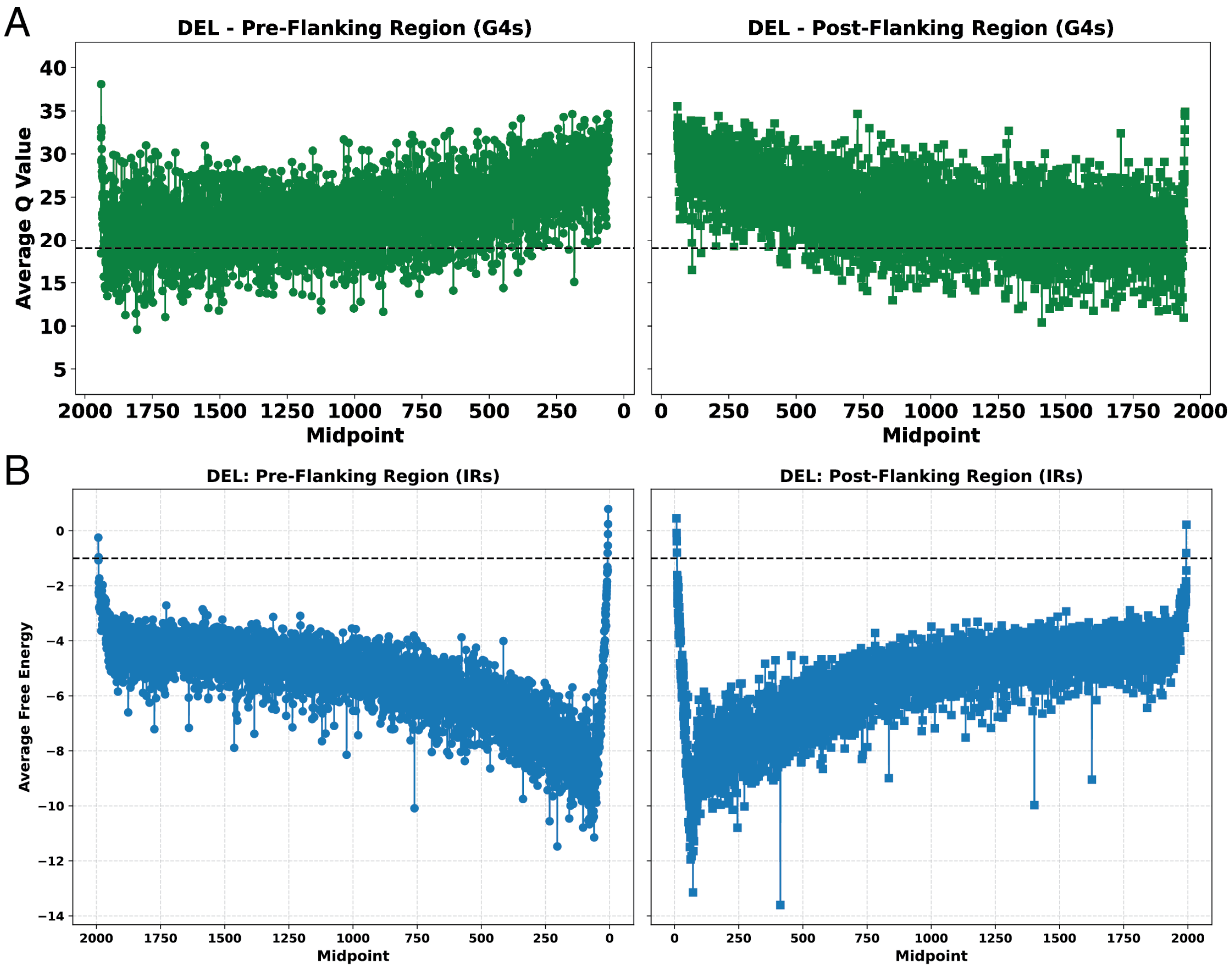


**Supplementary Figure 6 A-B. Stability profiles of non-B DNA motifs flanking deletion breakpoints. (A).** Average G4 stability scores (Quadron Q-values) across 2kp flanking regions upstream (pre-flank) and. Downstream (post-flank) of deletion (DEL) breakpoints. **(B).** Average free energy profiles of IRs across the same upstream and downstream deletion flanking regions. For each panel, motif stability values are plotted as a function of distance from the deletion breakpoint. Dashed horizontal liens indicate the stability thresholds used to classify motifs as stable (Q > 19 for G4s; free energy < 0 kcal/mol for IRs).

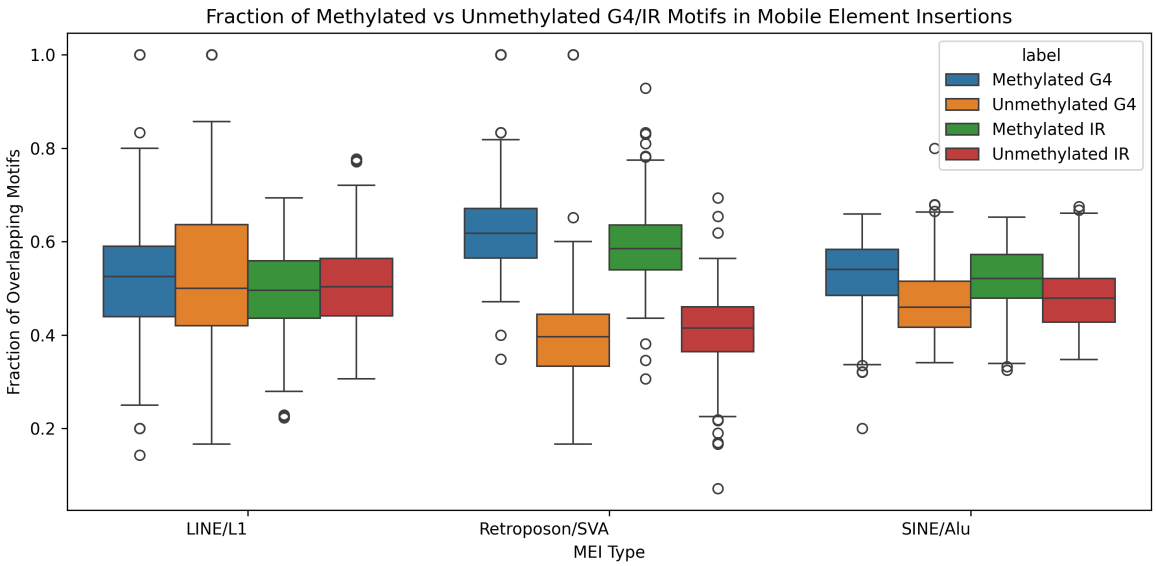


**Supplementary Figure 7. DNA methylation status of G-quadruplex and inverted repeat motifs within mobile element insertions.** Boxplots show the fraction of G4 and IR motif bases classified as methylation or unmethylated within major mobile element insertion families, including LINE/L1, Retroposon/SVA, and SINE/Alu elements. Motif-methylation overlap was assessed using Nanopolish-derived CpG methylation calls mapped to the GRCh38 reference genome, enabling integration of methylation information with mobile element annotations. Motif level methylation fractions are shown separately for methylated and unmethylated G4 and IR motifs within each mobile element insertion family.
